# Structural basis of dynein interaction with diverse activating adaptors

**DOI:** 10.64898/2026.04.25.720804

**Authors:** Ennio A. d’Amico, Sami Chaaban, Ferdos Abid Ali, Leon Michalski, Andrew P. Carter

**Affiliations:** Division of Structural Studies, MRC Laboratory of Molecular Biology, Cambridge, UK; School of Biochemistry, University of Bristol, Bristol, UK

## Abstract

Cargo-specific activating adaptors enable dynein to assemble with dynactin into processive supercomplexes. Adaptors share a coiled-coil architecture, but are highly diverse in sequence and structure, raising the question of how they converge on a common activation mechanism. To address this, we determined near atomic cryo-EM structures of dynein-dynactin assembled with five adaptors: RAB11FIP3, NIN, TRAK1, BICD2 and HOOK3. Despite their heterogeneity, all complexes contain adaptor coiled coils which bridge two dynein dimers to the dynactin filament. Adaptors are defined by an N-terminal interaction at the HBS1 with the dynein heavy chain, additional contacts along the dynein-dynactin groove, and C-terminal binding to the dynactin pointed end. However, we also found distinct sequence features, coiled-coil breaks and pointed-end interfaces that tune complex stoichiometry and stability. Our results define shared principles of dynein activation while revealing unexpected plasticity in how adaptors recognise and organise the dynein-dynactin machinery.

## Introduction

Cytoplasmic dynein-1 (dynein) is the main minus-end directed microtubule motor in human cells. It has a critical role in transporting and maintaining the spatial organisation of a broad range of cargoes, including endosomes, lysosomes, mitochondria, RNA granules, and signalling complexes^1^. Dynein recognises cargoes through a set of more than 20 identified and putative activating adaptors that provide the motor with cargo specificity^1,2^. All activating adaptors predominantly consist of coiled coils but vary in sequence and structure. While recent advances have shown examples of how dynein interacts with some adaptors^3–7^, the general principles for dynein recognition of an activating adaptor remain unclear.

At 1.4 MDa, dynein is among the largest cytoskeletal machines. Each motor is built of two heavy chains (DHCs), comprising the motor domain and the tail, two intermediate (DIC) and two light-intermediate chains (DLIC), and dimers of three light chains (LC8, Roadblock, Tctex)^8,9^. Dynein requires the co-factor dynactin to assemble a processive dynein-dynactin-adaptor (DDA) complex^8,10^. Dynactin is a ∼1.1 MDa assembly of 23 subunits and 11 distinct polypeptides^11^. It assembles around an actin-like minifilament of one actin and eight ARP1 subunits, capped by CapZα/β at the barbed end and by the four-subunit pointed-end complex (p25, p27, p62, ARP11) at the opposite end^11^. Within the DDA complex, the activating adaptor’s coiled coil positions dynein along the dynactin filament, determines the number of dynein motors recruited to a single supercomplex, and stabilises dynein into its activated form^3,8,11,12^.

In addition to coiled coils, activating adaptors contain an N-terminal domain which binds the dynein LIC subunit of dynein. Adaptors are grouped into distinct families based on the N-terminal LIC-binding domain they contain^1^. These include the CC1 box^15^, the HOOK domain^16^, EF-hands^17^, and the RH1 domain^4,18^. Most, although not all, known adaptors also contain a C-terminal Spindly motif (LΦXEΦ, with Φ being any hydrophobic residue) which binds the p25 subunit of dynactin^3,4,13,14^. In addition to the Spindly binding site on p25, referred to as site 4, previous work identified three other interfaces for adaptor binding on dynactin’s pointed end, distributed across p62 (sites 1-2) and p25 (site 3)^13^. Yet, the sequences different adaptors use to engage these sites are not known.

Between the LIC-binding module and the pointed end, adaptor coiled coils make additional contacts with the dynein heavy chains^11,12^. The Heavy chain Binding Site 1 (HBS1), has been structurally and functionally defined^3,6^. The HBS1 was first identified as the sequence QxxH/Y^6,7^, which contacts Tyr^827^ of dynein, and a nearby acidic patch which interacts with Arg^759^ on the same heavy chain^3^. This site is conserved among a subset of adaptors including BICDR1, HOOK3, TRAK1/2, and Spindly^3,6,7,19^. However, recent work shows that adaptors lacking this canonical sequence can still engage Tyr^827^ and Arg^759^ through different sequences^4,5,20^. Structural information is therefore necessary to understand how the currently uncharacterised adaptors recognise this key interaction site.

DDA complexes can be formed with one or two copies of the dynein dimer^11,12^. Stoichiometry in turn influences key motile parameters, including velocity, run length, and stall force^12,21^. While one-dynein complexes were identified first^11^, two-dynein complexes are more prevalent^3^, likely due to intrinsic properties of the interaction between the dynein tails and dynactin^22^. Adaptor identity biases the stoichiometry of the complex toward the presence of one or two dyneins both in cryo-EM structures off microtubules^12^, and in single-molecule fluorescence assays^12,23^. Additionally, adaptors themselves can be recruited singly or in pairs^3^. For many adaptors, the stoichiometry of the complexes they form is not known.

Here, using high-resolution structural analysis of complexes containing the adaptors RAB11FIP3, NIN, TRAK1, BICD2, and HOOK3, we address the structural principles underlying interactions between adaptors, dynein, and dynactin, and the features that enable adaptors lacking canonical motifs to still activate dynein motility.

### Organisation of DDA complexes

To solve DDA structures bound to microtubules, we improved previously-established image processing pipelines^3–5,24^, including the crucial step of subtracting the microtubule signal from integrated cryo-EM micrographs (Fig. s1). The output from pre-processing is a medium-resolution map of the adaptor-binding subcomplex (dynactin, adaptor, dynein tails including the heavy chains, motors, and part of the accessory chains), where the motors are less well-ordered due to flexibility but still visible at low threshold. We used the medium-resolution map for focussed classifications and refinements on sites where the adaptor contacted dynein and dynactin.

Using this pipeline, we determined cryo-EM structures of MT-bound DDAs assembled with adaptors belonging to different LIC-binding families: EF-hand (RAB11FIP3 and NIN), CC1-box (TRAK1 and BICD2), and Hook-domain (HOOK3) (Fig. 1a-g). To enhance complex formation, we used adaptor constructs that lack autoinhibition (Supplementary table 1). Unless otherwise indicated, we included the dynein co-factor LIS1, which enhances complex formation^23,25,26^, and a dynein motor point mutant which is less likely to form the autoinhibitory phi-particle^9^. In our comparative analyses, we also included previously reported structures of DDA complexes with BICDR1, JIP3, and NUMA1^3–5^ (Fig. 1h-j).

**Figure 1:**
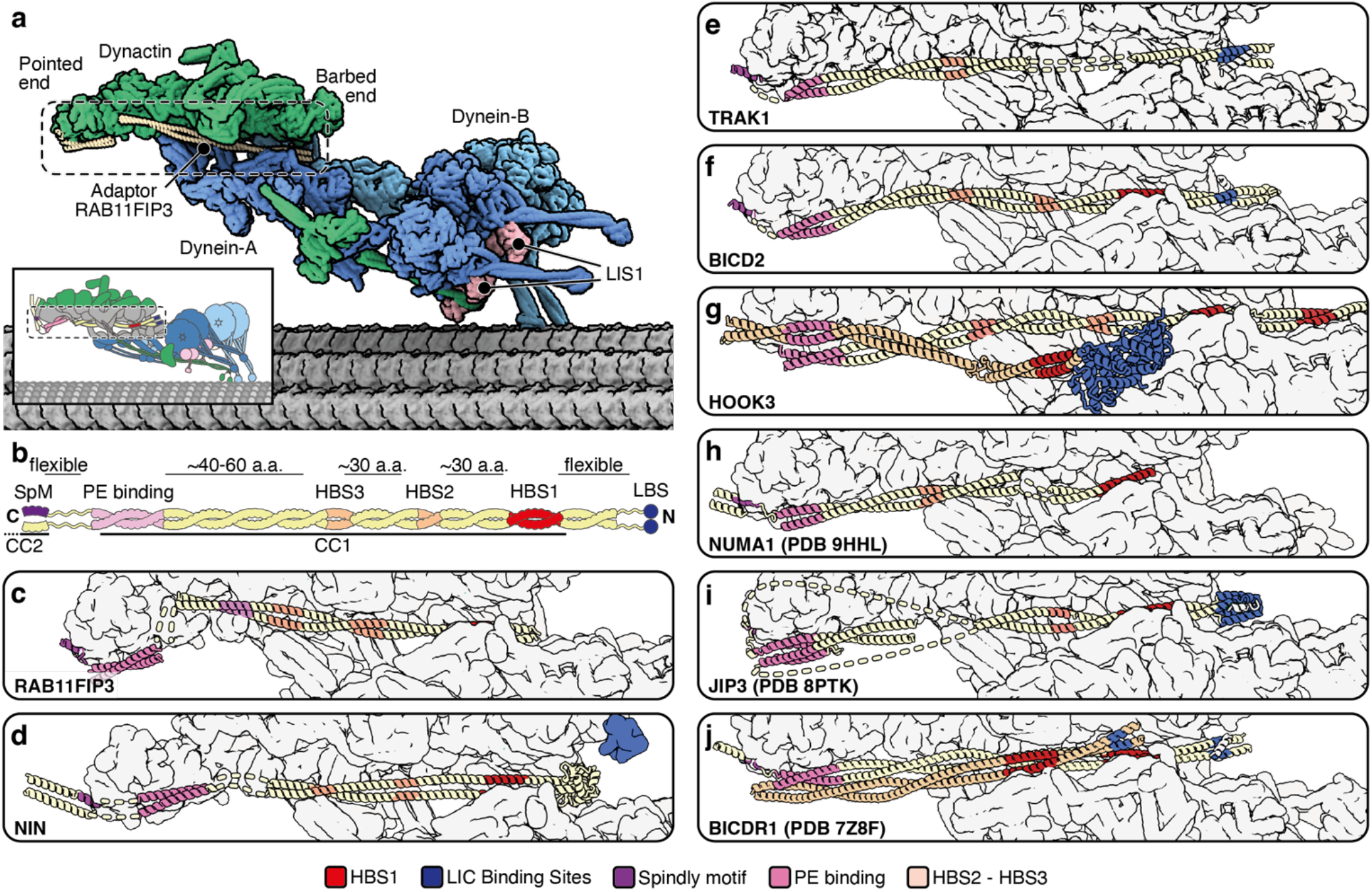
Architecture of DDA complexes. **a**, space-filling model of a DDA complex, with RAB11FIP3 as the adaptor, overlaid onto a 13-protofilament microtubule. **b**, schematic of an adaptor dynein-dynactin binding domain, with the main common contacts labelled. SpM, Spindly motif; PE binding, Pointed end-binding sites; HBS1-3, Heavychain-Binding Sites 1-3; LBS, Light intermediate chain-Binding Sites; a.a., amino acids **c-j**, DDA complex gallery. Only the core region of the complex, which includes the main interaction points between dynein-dynactin and the adaptor is shown (dashed box in **a**).

Despite the diversity in domain organisation, coiled coil length, and sequence, all adaptors use a limited number of sites to interact with dynein and dynactin. Thus, the overall architectures of all the DDA supercomplexes are similar to each other and to those reported previously (Fig. 1b, Fig. 1c-j). All adaptors in our study can recruit two dyneins. Following established nomenclature, the dynein closest to the pointed end is termed dynein-A, while the dynein nearer to the barbed end is termed dynein-B, each made of two copies (A1, A2 and B1, B2 respectively^3,12^; Fig. 1a). The only adaptor where we observed a significant class of supercomplex containing one dynein was BICD2^12^. Furthermore, only HOOK3 and BICDR1 DDAs have the propensity to recruit two copies of the adaptor^3^ (Fig.1g, j). In both cases one adaptor binds canonically, placed in the groove between dynactin and dynein-A, and the second copy sits on top of the first, in the case of HOOK3, or below, in the case of BICDR1, as previously reported^3^. We do not see in either case a class of complexes with only a single adaptor bound.

The canonical adaptor binding mode has the N-terminal coiled coil of the adaptor bridging the cleft between the dynein tails and the dynactin filament, with the adaptor’s N-terminus oriented towards the barbed end of dynactin and its C-terminus towards the pointed end. N-terminal to the coiled coil, the LIC-binding module and LIC interaction are only clearly resolved in CC1-box adaptors TRAK1 and BICD2 (Fig. 1e-f).

Within the coiled coil regions, the adaptors are most ordered close to dynactin’s barbed end due to the HBS1 interaction. Most adaptors also interact with the tails of dynein-A1 and A2 after the HBS1. Here, we therefore refer to these interactions as HBS2 and HBS3 (Fig. 1b). The coiled coil stability in this region is variable, with some adaptors displaying a continuous coiled coil and others displaying breaks in variable positions. Following established nomenclature, we call the coiled coil which bridges dynein and dynactin CC1, or CC1a and CC1b in case of breaks^5,7,27^.

Most adaptors make two contacts with dynactin’s pointed end complex. There are Spindly motifs in all adaptors we analysed^15^. These lie within loops after the coiled-coil break which ends CC1, or at the beginning of a CC2 coiled-coil segment. In addition, most adaptors bind the pointed end at sites 2 and/or 3 via the C-terminus of their CC1. The only exception is JIP3, where two coiled coil fragments C-terminal to the Spindly motif make the pointed end interaction (Fig. 1i).

In the following sections we will use our high-resolution structures to describe the molecular basis for the way five different adaptors recruit dynein-dynactin.

### Structure of the Dynein-Dynactin-RAB11FIP3 complex

RAB11FIP3, also known as FIP3, is an effector of the small GTPase RAB11A (Fig. 2a)^28^. It has critical roles in the integrity of the endosomal recycling compartment^28^, and in ciliary targeting of receptors like rhodopsin^29^. FIP3 mutations can cause defects in the cell cycle by impairing cytokinesis^30^. FIP3 activates dynein processivity *in vitro*^10^ and interacts with the dynein LIC through its N-terminal EF-hand domains in a calcium-independent manner^17^.

**Figure 2:**
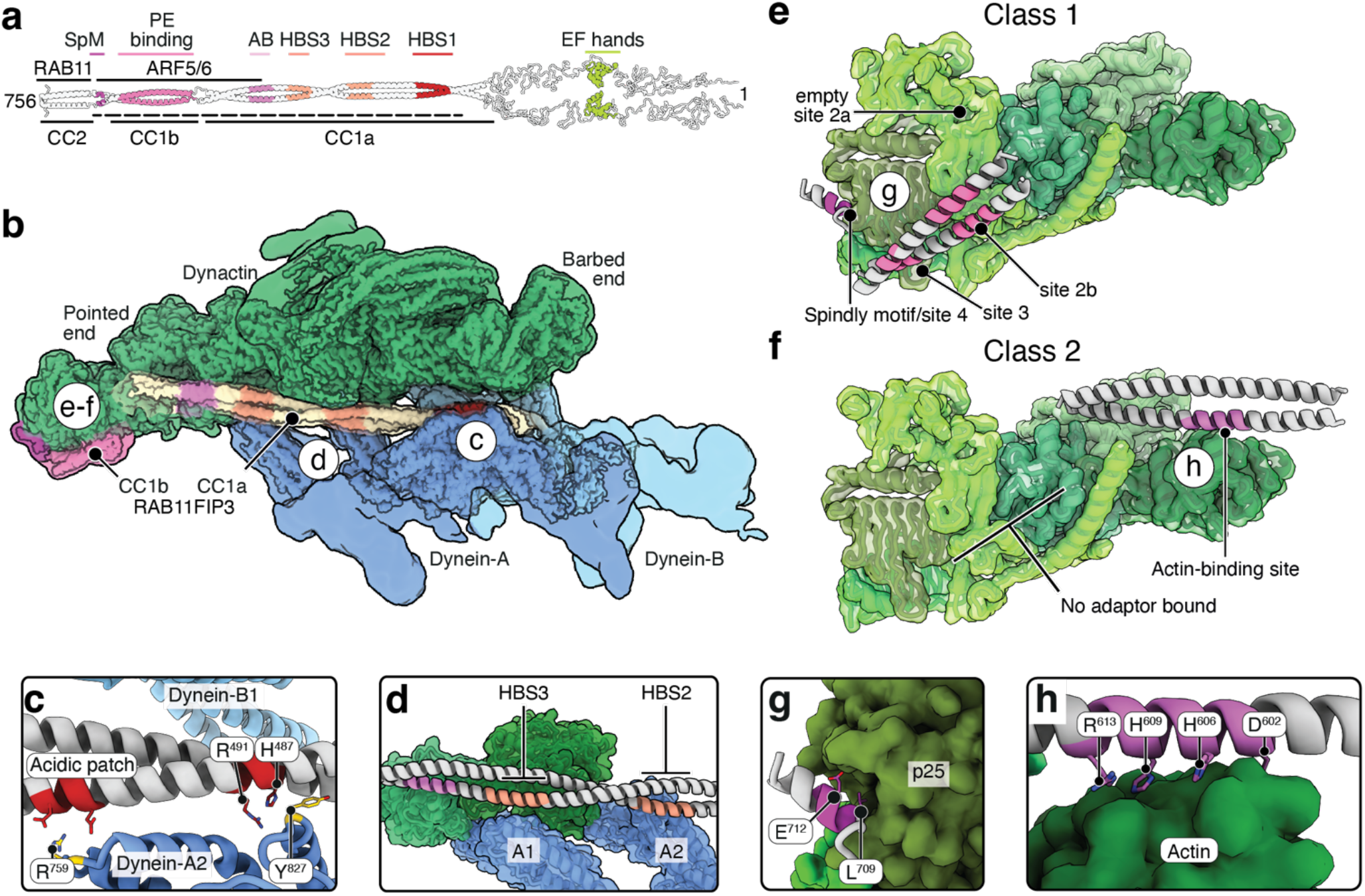
Structure of the dynein-dynactin-FIP3 complex. **a**, AF2 prediction of full-length FIP3 with coloured domains for interaction with dynein and dynactin. Dashed line marks part of the adaptor which is visible in our experimental density. SpM, Spindly motif. PE binding, pointed end-binding site. AB, actin binding site. HBS1-3, heavy chain-binding site 1-3. ARF6 and RAB11 binding sites in black according to published literature. **b**, Composite density map for the dynein-dynactin-RAB11FIP3 complex. The dynein motors are flexible relative to the adaptor-interaction module and are therefore not resolved here. **c**, Interaction of FIP3 with the dynein heavy chains at the HBS1. **d**, organisation of HBS2/3 contacts of FIP3 with the tails of dynein-A. **e**, organisation of contacts between FIP3 and the dynactin pointed end (class 1). **f**, organisation of RAB11FIP3 contacts with the actin subunit of dynactin (class 2). **g**, Interaction between p25 and the FIP3 Spindly motif. **h**, Interaction between actin and FIP3.

In our structure of dynein-dynactin-FIP3-LIS1, FIP3 shows a break in the CC1 coiled coil, with CC1a occupying the canonical cleft between two dynein dimers and dynactin and CC1b binding to the pointed end (Fig. 2b; Fig. s2a-b). We were unable to visualise the EF-hands, which are connected to the adaptor coiled coil by a long unstructured loop and likely flexibly positioned. The HBS1 of FIP3 lacks the conserved QxxH/Y motif^6^. Instead, the adaptor’s His^487^ interacts with dynein-A2’s Tyr^827^ and the nearby Arg^491^ contacts the backbone of the dynein heavy chain (Fig. 2c; Fig. s2c). A conserved acidic patch of residues Glu^506^ and Glu^510^ on FIP3 interacts with dynein’s Arg^759^ (Fig. 2c). The HBS2 and HBS3 interactions involve two charged patches on FIP3, residues 535-550 and 575-590, which bind to the tails of dynein-A2 and A1, respectively (Fig. 2d).

FIP3 binds dynactin atypically in two mutually exclusive modes which are both present in our sample (Fig. 2e-f). In class 1, the adaptor contacts the dynactin pointed end through CC1b and its Spindly motif, but the C-terminal segment of CC1a is disordered (Fig. s2d). The CC1b engages with the bottom part of site 2^13^, which we now call site 2b, and not with the top part (site 2a) (Fig. 2e, Fig. s2d). The resolution of our maps is sufficient to assign the Spindly motif to residues 709-713, corroborated by sequence conservation and AlphaFold2 (AF2) prediction (LAAEI; Fig. 2g). In class 2, the CC1a binds the actin subunit of the dynactin filament, but the adaptor is disengaged from both pointed end contacts (Fig. 2f, Fig. s2d). The FIP3 side chains involved with contacting actin are Asp^602^, His^606^, His^609^ and Arg^613^ (Fig. 2h).

Like all adaptors, FIP3 interacts with cargo through its C-terminal domain, which is atypically close to the dynein-binding region and even overlaps with it^31,32^. The adaptor binds RAB11A through a short site, C-terminal to the Spindly motif^31,32^. It also binds small GTPases ARF5 and ARF6 through a domain located between residues 607-712^32^, mapping to the end of CC1a all of CC1b, and Spindly motif (Fig. 2a). Therefore, RAB11A recruitment is compatible with full engagement of FIP3 with dynactin, but ARF5/6 recruitment is not.

Together, these observations indicate that FIP3 alternates between two distinct anchoring modes, potentially reflecting a mechanism to modulate access to the cargo binding site and thereby coordinate cargo attachment with dynactin recruitment.

### Structure of the Dynein-Dynactin-NIN complex

NIN (also known as Ninein) and its paralog NINL (Ninein-like) localise at the centrosome and are involved in microtubule nucleation^33,34^. NINL has also been implicated in melanosome transport^35^ and in responding to interferon during viral infection^36^. While both proteins are large (NIN, 2090 a.a., NINL, 1382 a.a.), only the EF-hands, CC1 and part of CC2 are necessary to activate dynein motility^34^. Both proteins contain two pairs of EF-hand domains. In NIN, only EF-hand 1 (EF-1) mediates the interaction with the LIC^17^. The EF-hand pairs are followed by pseudo-EF hands immediately before the start of the coiled coil (Fig. 3a). No HBS1 or Spindly motifs could be identified prior to this work in either NIN or NINL.

**Figure 3:**
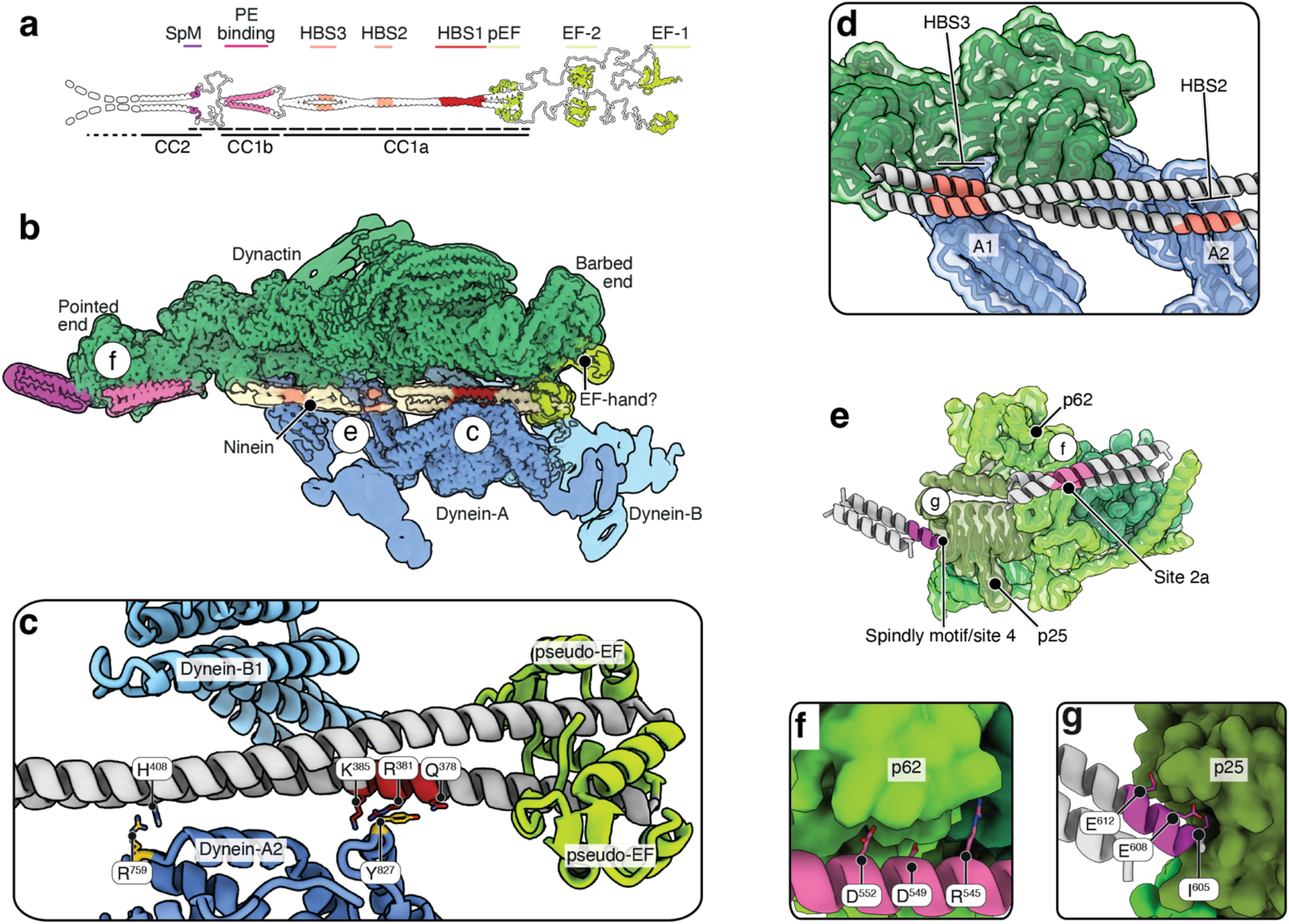
Structure of the dynein-dynactin-NIN complex. **a**, AF2 prediction of dynein-binding region of NIN with coloured domains for interaction with dynein and dynactin. Dashed line marks part of the adaptor which is visible in our experimental density. SpM, Spindly motif. PE binding, Pointed end-binding site. HBS1-3, heavy chain-binding site 1-3. pEF, pseudo-EF hand pair. EF, EF-hand pair. **b**, Composite density map for the dynein-dynactin-NIN complex. The dynein motors are flexible relative to the adaptor-interaction module and are therefore not resolved here. **c**, structure of the NIN interaction with the dynein HBS1. **d**, organisation of HBS2/3 contacts of NIN with the tails of dynein-A. **e**, density map of the dynactin pointed end and NIN, with the different subunits and motifs coloured. **f**, structure of the interaction between NIN and p62 on dynactin. **g**, structure of the NIN Spindly motif interaction.

The NIN CC1a follows the groove between dynein and dynactin from its start immediately after the pseudo-EF-hand domain (Fig. 3b; Fig. s3a-b). Extra density is present adjacent to the barbed end of dynactin, which matches the size of an EF-hand pair, but has very low resolution, preventing confident assignment (Fig. s3c). The NIN HBS1 involves Gln^378^ and Arg^381^ making contacts with dynein-A2 Tyr^827^ (Fig. 3c, Fig. s3d). NIN also interacts with the backbone of the A2 heavy chain through Lys^385^. Uniquely among currently known adaptor HBS1s, NIN lacks an acidic patch. Instead, His^408^ binds dynein Arg^759^ (Fig. 3c; Fig. s3d). Within the groove, downstream of the HBS1, the NIN density is smeared (Fig. 3b), indicating a high degree of flexibility. Nonetheless, we could identify HBS2 and HBS3 contacts with the tails of dynein A2 and A1, through residues 440-450 and 480-490, respectively (Fig. 3d), although the density is weaker than at the HBS1.

NIN interacts with the dynactin pointed end subunits p62 and p25 (Fig. 3e; Fig. s3e). The interaction with p62 at site 2a is well-resolved and involves the adaptor residues Arg^545^, Asp^549^, Asp^552^ (Fig. 3f; Fig. s3e). The interaction with p25 at site 4 uses a Spindly motif of sequence IETEL (residues 605-609) (Fig. 3g, Fig. s3e). An extra point of interaction at Glu^612^ (+8 from the start of the Spindly motif) contributes to stabilising the adaptor further and limits its flexibility close to the pointed end (Fig. 3g). This may explain why the coiled coil that extends C-terminally from the Spindly motif is resolved further than other adaptors investigated here (Fig. 3b, Fig. s3e). Thus, NIN binds dynein-dynactin through an atypical HBS1 and a Spindly motif that differs from the LΦXEΦ consensus.

### Structure of the Dynein-Dynactin-TRAK1 complex

TRAK1 links mitochondria to the microtubule-based motor machinery^37^. It binds both kinesin-1 and dynein through an N-terminal coiled coil, and localises to neuronal axons^38^. TRAK1 contains a CC1 box motif (AAxxG), a conventional HBS1 containing a QxxH motif and an acidic patch, and a Spindly motif^19^. Uniquely, it displays an extremely short N-terminal coiled coil (CC1a) which harbours both the CC1 box and the HBS1, followed by a coiled coil break starting within the acidic patch (Fig. 4a). Unlike most other adaptors, the density for TRAK1 is unresolved within the dynein groove between the HBS1 and HBS3 contacts (Fig. 4b-c; Fig. s4a-b). The unresolved fragment consists of a large part of CC1b, and contains the mapped kinesin-1-binding region (residues 110-270)^19,39^. At low threshold, some extra density is present next to the A2 tail, suggesting a very weak HBS2 contact (Fig. s4c).

**Figure 4:**
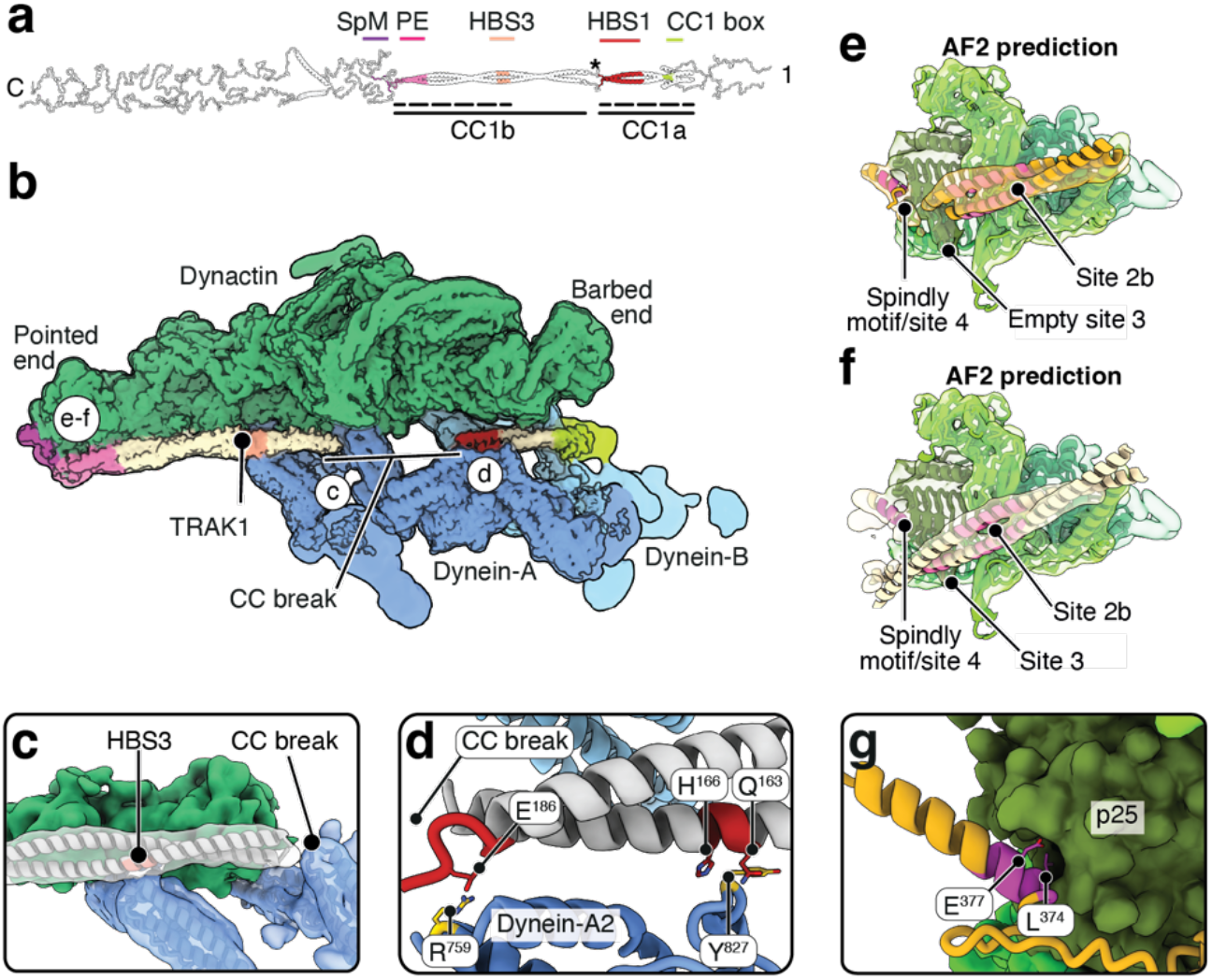
Structure of the dynein-dynactin-TRAK1 complex. **a**, AF2 prediction of full length TRAK1 with coloured domains for interaction with dynein and dynactin. Stars mark coiled-coil breaks. Dashed line marks part of the adaptor which is visible in our experimental density. SpM, Spindly motif. PE binding, Pointed end-binding site. HBS1-3, heavy chain-binding site 1-3. **b**, Density map for the dynein-dynactin-TRAK1 complex. The dynein motors are flexible relative to the adaptor-interaction module and are therefore not resolved here. **c**, Model of HBS3 contacts between TRAK1 and Dynein-A1, fit into a filtered density map for the HBS2/3 region of the DDT complex. **d**, AF2 prediction of the interaction between dynein-A2 heavy chain and the HBS1 on TRAK1. **e**, Organisation of the contacts of TRAK1 with the dynactin pointed end complex in class 1 (AF2 prediction), fit into class 1 density. The p62 saddle interaction is occupied, but site 3 on p25 is not. Residues with pLDDT < 60 or far from the pointed end complex hidden. **f**, Organisation of the contacts of TRAK1 with the dynactin pointed end complex in class 2 (AF2 prediction), fit into class 2 density. Residues with pLDDT < 60 hidden. **g**, Structure of the interaction of the Spindly motif on TRAK1 with the p25 subunit of dynactin (AF2 prediction for class 1 interaction).

Within CC1a, we can resolve both the CC1 box and the HBS1 of TRAK1, albeit to lower resolution than other adaptors. In AF2 predictions, the CC1 box of TRAK1 is shielded by autoinhibitory helices^39^ (Fig. 4a). We are unable to distinguish whether the LIC displaces the autoinhibitory helices in our structures, due to resolution limitations (Fig. s4c-d). However, given that the LIC-CC1 box interaction is necessary for function, we model the LIC as bound. The QxxH motif of the HBS1 (residues 163-166) interacts with Tyr^827^ and is well resolved. The acidic patch (residues 183-188) is more poorly resolved and overlaps with the end of CC1a (Fig. s4e). As expected, it contacts Arg^759^ on the dynein heavy chain.

TRAK1 contacts the pointed end through sites 2b-3 and site 4 (Fig. 4e-f). Classification shows it either contacts site 2b (class 1) or site 2b and site 3 (class 2) which matches two alternative AF2 predictions, in which the adaptor binding is offset by ∼25 residues (Fig. 4e-f; Fig. s4f). At site 4, the adaptor is bound similarly in both classes, with an interaction mediated by the Spindly motif (LAAEI; residues 374-378; Fig. 4g). Overall, the TRAK1 coiled coil domain interacts with dynein with unusual flexibility, which allows for two broadly different interactions with the dynactin pointed end and may be important for dynein and kinesin-1 to bind TRAK1 at the same time.

### Structure of the Dynein-Dynactin-BICD2 complex

BICD2 (Fig. 5a) recruits dynein to Rab6-positive secretory and Golgi-derived vesicles^40^, supporting their minus-end-directed transport and Golgi organisation in many cell types. Mutations in BICD2 are associated with dominant spinal muscular atrophy^41^, highlighting its requirement for dynein-mediated transport in neuronal function. More recently, BICD2 has been shown to be exploited by pathogens, like HPV^42^ and *O. tsutsugamushi*^43^. Unlike BICDR1 and HOOK3, structures of the dynein-dynactin-BICD2 complex, solved without microtubules, included a mixture of one-dynein and two-dynein particles^9,11^. On microtubules, data was more contradictory: while single-molecule fluorescence microscopy data indicated that BICD2 has a higher propensity to form complexes which include only a single dynein dimer^12,23^, early tomography studies indicated an overwhelming prevalence of two-dynein complexes^44^. Furthermore, the lack of side chain resolution for the adaptor prevented identification of the main binding sites. Previous literature identified a putative conserved QxH HBS1 in BICD2^6^, which resembles only partially the QxxH/Y motif.

**Figure 5:**
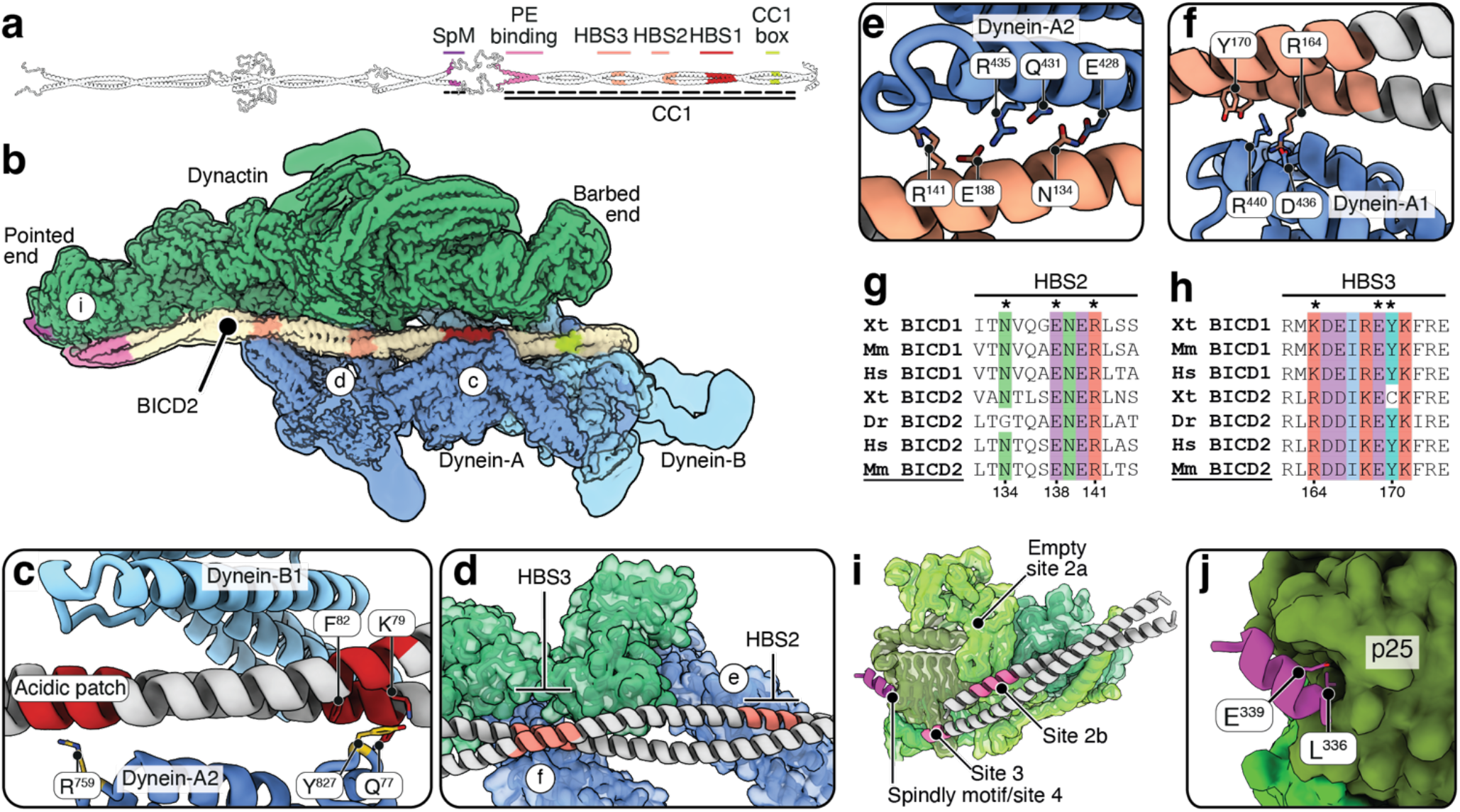
Structure of the dynein-dynactin-BICD2 complex. **a**, AF2 prediction of full length BICD2 with coloured domains for interaction with dynein and dynactin. Stars mark coiled-coil breaks. Dashed line marks part of the adaptor which is visible in our experimental density. SpM, Spindly motif. PE binding, Pointed end-binding site. HBS1-3, heavy chain-binding site 1-3. **b**, Composite density map for the dynein-dynactin-BICD2 complex when two dynein dimers are present. The dynein motors are flexible relative to the adaptor-interaction module and are therefore not resolved here. **c**, Structure of the interaction between dynein-A2 heavy chain and the HBS1 on BICD2. **d**, Organisation of HBS2 and HBS3 contacts between BICD2 and dynein-A. **e**, Structure of the contacts between dynein-A2 heavy chain and the HBS2 on BICD2. **f**, Structure of the accessory contacts between dynein-A1 heavy chain and the HBS3 on BICD2. **g**, Sequence alignments in the BICD2 family for the HBS2. **h**, Sequence alignments in the BICD2 family for the HBS3. **i**, Organisation of adaptor position on the dynactin pointed end. **j**, Structure of the Spindly motif of BICD2 bound to the pointed end.

In the BICD2 dataset, unlike all other adaptors, around 20% of particles lack dynein-B and form a dynein-A only complex (Fig. 5b, Fig. s5a-c). Conversely, we do not detect complexes where only dynein-B is present. One-dynein complexes exhibit significantly worse density for dynein-A compared to two-dynein complexes, suggesting increased flexibility (Fig. s5c). To understand the cause, we compared the position of dynein-A in one vs two-dynein complexes (Fig. s5d). The interactions between dynein-A and the adaptor are similar to those in the two-dynein complex. However, upon binding of the second dynein, contacts between dynein-A and dynein-B cause the adaptor and the dynein-A tails to swing downwards and backwards as a rigid body relative to dynactin (Fig. s5d). These movements are consistent with what was observed in previous studies off microtubules^12^. The interactions between dynein-B and dynein-A^3^ contribute to stabilising the complex and reducing the flexibility of the dynein-A tails. The observations on the specific interactions between BICD2 and dynein-dynactin described below are based on the two-dynein structure but also apply to the one-dynein complex we visualise.

BICD2 interacts with the dynein LIC through a CC1 box^16^. We could resolve only a single copy of the LIC-binding helix, on the face of the adaptor which is closest to dynein-B2 (Fig. s5e). Our structure shows that the HBS1 interaction between dynein and BICD2 is not mediated by the proposed QxH motif. Instead, the adaptor residues Lys^79^ and Phe^82^ from one chain and Gln^77^ of the opposite chain contact dynein’s Tyr^827^ (Fig. 5c; Fig. s5f). A conserved acidic patch (residues 101-106) interacts with dynein’s Arg^759^ (Fig. 5c). BICD2 makes HBS2 and HBS3 contacts with previously identified sites on the tails of dynein A2 and A1, respectively^11^ (Fig. 5e-h). The interaction with dynein-A2 is mediated by charge-complementarity (Fig. 5e, g), and with dynein-A1 by a highly conserved charged patch including residues Arg^171^, Asp^175^, and Tyr^176^ (Fig. 5f, h). The latter contacts involve both BICD2 chains. The adaptor binds the dynactin pointed end through the terminal part of CC1 and the beginning of CC2, respectively interacting with sites 2b-3 and 4 (Fig. 5i-j). The contact with site 2b-3 is flexible and we are unable to resolve side chains. After a long unresolved loop, binding to site 4 is mediated by a conserved Spindly motif^15^ (LLSEL, residues 336-340; Fig. 5j).

### Structure of the Dynein-Dynactin-HOOK3 complex

HOOK3 is a member of the highly conserved Hook domain family of adaptors, present through evolution to filamentous fungi like *A. nidulans*^45^.

Mammalian HOOK proteins are a component of the tripartite Fts-HOOK-FHIP complex which activates and links dynein to mostly endosomal and Golgi cargoes^46,47^. HOOK3 binds the LIC through the Hook domain^17^, displays a conventional HBS1 with a QxxH motif and acidic patch^3,6^, and a conserved Spindly motif^5^ (Fig. 6a). Our previous studies showed two HOOK3 adaptor dimers forming part of a single DDH motile complex^3,12^. However, these structures were solved at low resolution and off microtubules, preventing correct assignment of the registry of the coiled coil and identification of interacting residues.

**Figure 6:**
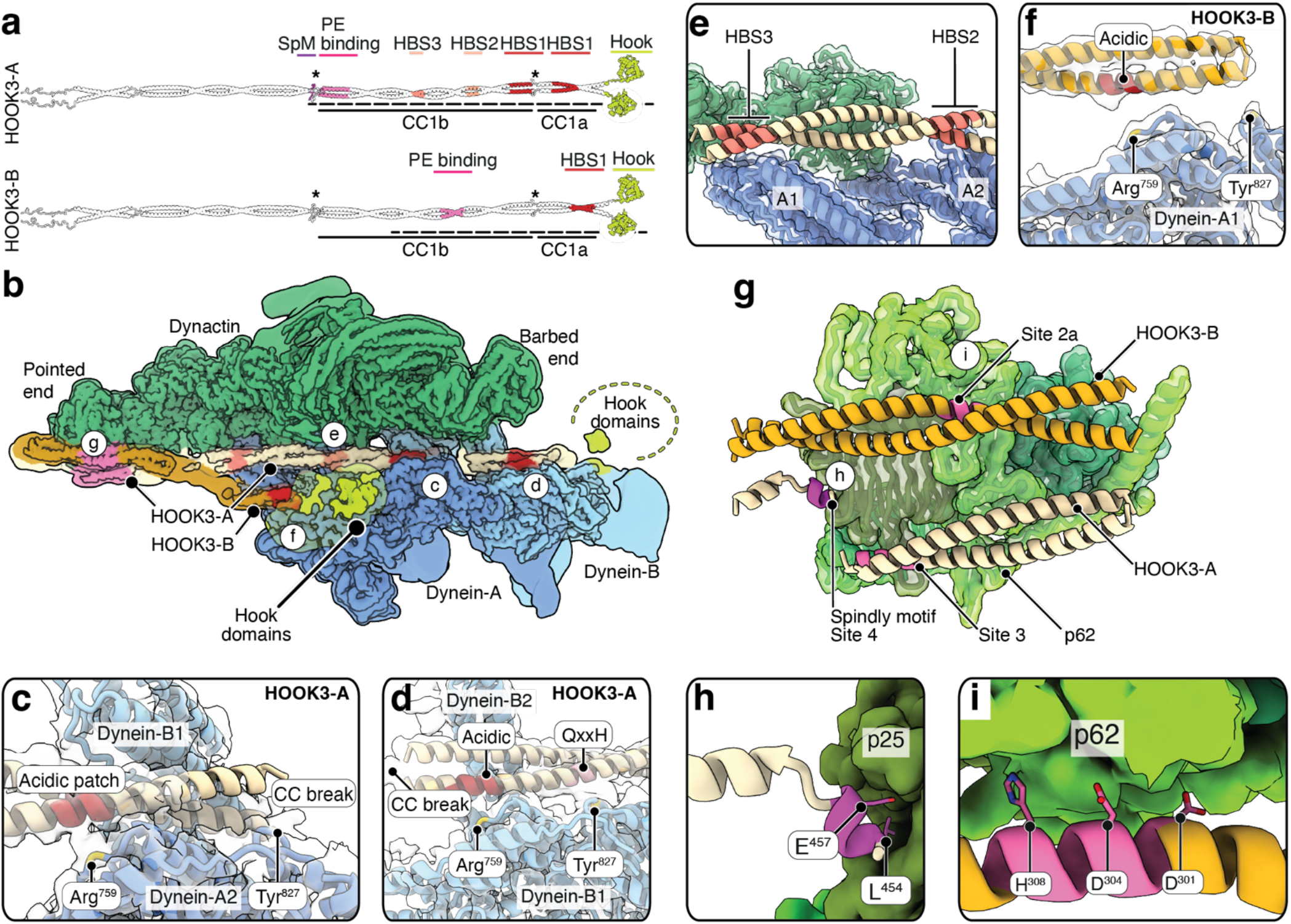
Structure of the dynein-dynactin-HOOK3 complex. **a**, AF2 prediction of full length HOOK3 with coloured domains for interaction with dynein and dynactin. Stars mark coiled-coil breaks. Dashed line marks part of the adaptor which is visible in our experimental density. SpM, Spindly motif. PE binding, Pointed end-binding site. HBS1-3, heavy chain-binding site 1-3. **b**, Composite density map for the dynein-dynactin-HOOK3 complex. The dynein motors are flexible relative to the adaptor-interaction module and are therefore not resolved here. **c**, Model for the interaction at the HBS1 between HOOK3-A, dynein-A2 and the tail of dynein-B1. **d**, Model for the interactions between HOOK3 and the tails of dynein-A2 and A1 at the HBS2 and HBS3. **e**, Interaction between dynein-B1 and HOOK3-A at the secondary HBS1. **f**, Interaction between dynein-A1 and HOOK3-B at the HBS1. **g**, Interaction between both HOOK3 dimers and the dynactin pointed end. **h**, Interaction between HOOK3-A and the dynactin pointed end at the Spindly motif. **i**, Interaction between HOOK3-B and p62 at site 2.

In contrast to our previous work^3,12^, in this new structure, all the interactions of both adaptors with dynein and dynactin are visible (Fig. 6a-b, Fig. s6a-b). The main adaptor (HOOK3-A) is resolved from its N-terminal Hook domain, through its short CC1a, a longer CC1b, to the C-terminal Spindly motif. Like all other adaptors, HOOK3-A binds the dynein heavy chain of dynein-A2. Unusually, this involves an HBS1 at the very start of its CC1b segment, rather than CC1a. Unlike all other adaptors in this study, there is no contact with dynein Tyr^827^, and the segment of adaptor closest to this site is highly flexible (Fig. 6c). There is however a conserved acidic patch centred around residues Asp^266^ and Asp^267^ which binds to dynein’s Arg^759^ (Fig. 6c). Also unique is that HOOK3-A makes an additional contact with dynein-B1 through its CC1a (Fig. 6b, d; Fig. s6d). Previous studies had identified a QxxH conserved motif on HOOK3^6^ and AlphaFold2 predicts an HBS1 interaction with dynein through this site^3^ and an acidic patch downstream of it, around Glu^219^ (Fig. 6d). In our structure, this prediction matches the interaction between HOOK3-A and dynein-B1 rather than the usual dynein-A2. Therefore, HOOK3-A uniquely contains two HBS1s (Fig. 6a), after which an HBS2 and HBS3 form charged interactions with dynein-A2 and dynein-A1 respectively (Fig. 6e).

The second adaptor (HOOK3-B) is resolved from the N-terminal Hook domain until around halfway along CC1b (Fig. 6a). The globular density for the Hook domain is present close to dynein-A1, which allows us to define that CC1a contacts this chain in an HBS1-like manner (Fig. s6e). While the resolution is not sufficient to resolve side chains, we are able to use the globular density of the Hook domain and coiled-coil breaks to place the CC1a (Fig. 6f). This suggests that the HBS1 on HOOK3-B is different from the two HBS1s on HOOK3-A (Fig. s6f): it uses an acidic patch centred around residues Glu^193^ and Glu^194^ to contact Arg^759^ and does not contact Tyr^827^ (Fig. 6f). Therefore, HOOK3 recognises the dynein heavy chain through multiple HBS1 sequences.

The interaction of both HOOK3 adaptors with the dynactin pointed end is well resolved in our structure (Fig. 6g; Fig. s6g). HOOK3-A binds site 3 via the end of CC1b and, following a short loop, to site 4 through its Spindly motif (LAAEI, residues 454-458) (Fig. 6h). HOOK3-B is located above HOOK3-A and interacts with p62 through a lengthy interface which spans two main hotspots. Adaptor residues Asp^301^, Asp^304^ and His^308^ bind to site 2a similarly to NIN (Fig. 6i; Fig. s6g). Downstream of this, we detected a novel interface with a flexible loop within p62, close to site 2a. This segment is not normally resolved in other adaptors but is stabilised by the interaction with the HOOK3B (Fig. s6g). These results match and explain previously published crosslinking data which indicate that HOOK3 binds both sites 2 and 3^13^. The two adaptors are offset by roughly 110 residues on the pointed end, and the interfaces used for interaction at the pointed end markedly differ. Overall, our structure shows that HOOK3-A and HOOK3-B use independent sites to recognise the dynactin pointed end and dynein.

## Discussion

Our cryo-EM analysis of five activating adaptors, combined with previous work done on BICDR1, JIP3, and NUMA1^3–5^, reveals that dynein-dynactin-adaptor (DDA) assemblies share a conserved architecture, yet contain distinct, adaptor-specific interactions (Fig. 7a). The dynein-dynactin binding domain starts with one of four N-terminal LIC-binding domains. The core of the interaction between adaptors and dynein-dynactin is a coiled-coil domain (CC1), which can be continuous (BICD2, BICDR1) or broken (FIP3, NIN, TRAK1, NUMA1, JIP3, HOOK3). All adaptor CC1s contact the heavy chain of dynein-A2 via an HBS1. Most adaptors make further contacts with the tails of dynein-A2 and dynein-A1, which we name HBS2 and HBS3. The C-terminal end of CC1 contacts the dynactin pointed end at sites 2 or 3. A flexible linker separates CC1 from the Spindly motif, which binds p25 at site 4, and ends the dynein-dynactin binding domain.

**Figure 7:**
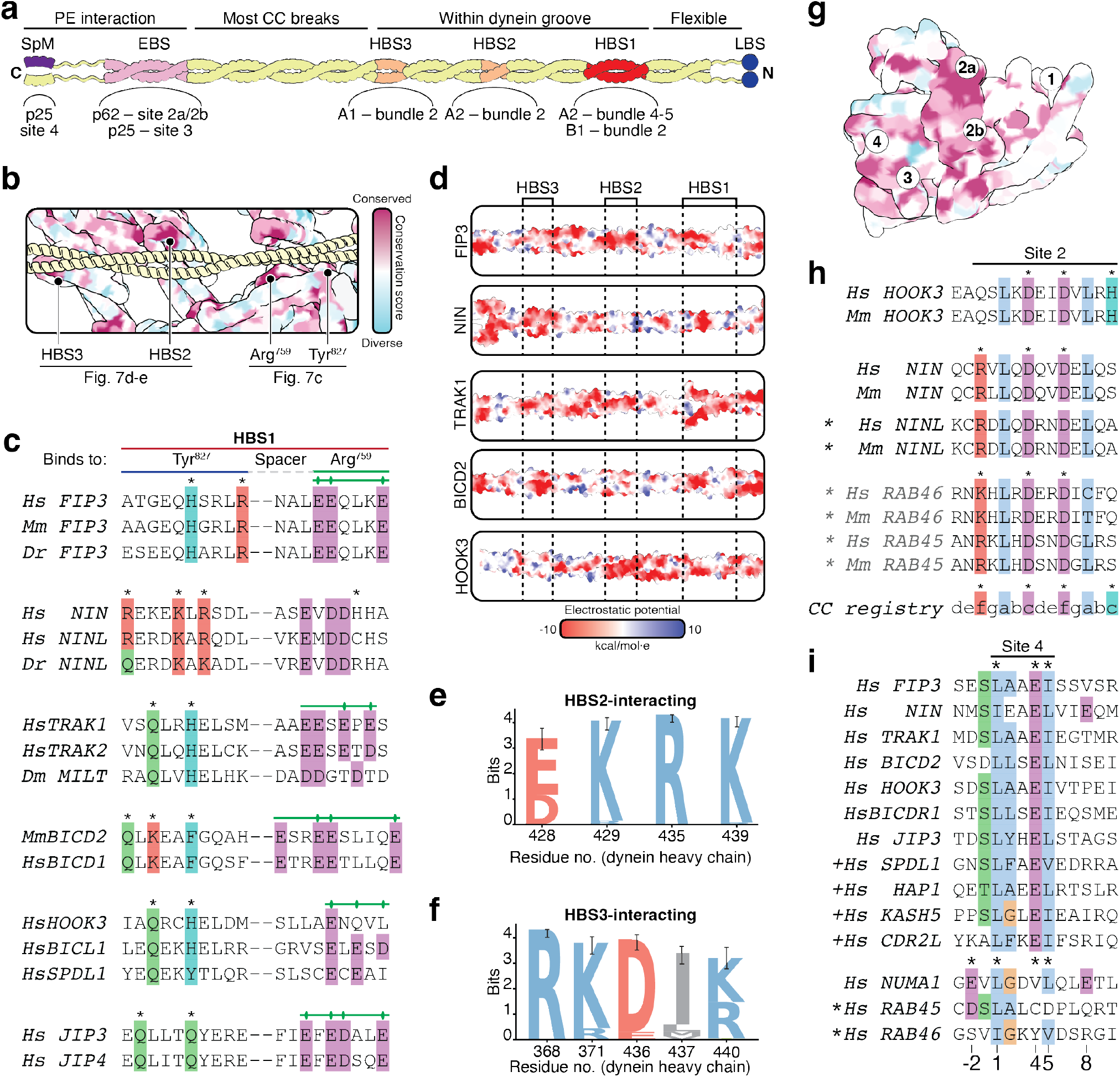
Shared features of dynein recognition of adaptors. **a**, Schematic of an adaptor with relevant binding sites displayed, and which region of dynein-dynactin they bind. **b**, Model of the dynein groove, coloured by conservation across metazoa, with a sample adaptor (BICD2) displayed. Core binding sites are labelled. **c**, Sequence conservation at the HBS1 in all adaptors screened and relevant homologues. **d**, Space-filling models for AF2 prediction of adaptors in this study, coloured by coulombic charge, indicate regular negatively charged patches. **e-f**, WebLogos of the conservation of residues that interact with the HBS2 (**e**) and HBS3 (**f**) among diverse metazoa (n = 52 sequences). Residues are coloured by chemical properties. Error bars represent 95% Wilson confidence intervals. **g**, Model of the dynein pointed end, coloured by conservation. Adaptor binding sites are labelled. **h**, Sequence alignment of adaptors binding to p62 through a site 2 interaction, showcasing the conserved organisation of charged residues. * sign indicates adaptors where the site 2 interaction was identified based on AF2 prediction. **i**, Sequence alignment of the Spindly motif across adaptor families. + sign indicates adaptors where the Spindly motif was identified based on sequence prediction. * sign indicates adaptors where a Spindly-like motif was identified based on AF2 prediction only.

Within this shared scaffold, adaptors diversify in three ways: the identity and spacing of dynein-facing contacts, whether and how the coiled coil is interrupted by breaks, and the geometry and multiplicity of interactions at sites 2 and 3. These parameters might tune stoichiometry and could be expected to contribute to complex stability and ultimately motile output.

### Dynein accommodates adaptor coiled coils with plasticity

The tail of a dynein heavy chain consists of a series of helical bundles^9^. The HBS1 of all adaptors bind the same region on the dynein-A2 heavy chain, at the interface between helical bundles 4 and 5 (Arg^759^ and Tyr^827^, respectively) ^3,6,7^. The adaptor’s HBS2 and HBS3, in contrast, contact helical bundle 2 in the tail of dynein-A2 and dynein-A1 respectively. On dynein-B1, helical bundle 2 binds to the other side of HBS1. The sites on dynein which recognise HBS1-3 are highly conserved across metazoa (Fig. 7b) but not in unicellular organisms like *A. nidulans* where adaptor-mediated transport over long distances is well-established^45,48^. The functional implications of this divergence remain unclear. Overall, these results suggest that adaptors might be recognised differently in unicellular organisms and metazoa.

While we identify HBS1 interactions in all adaptors, only a subset – most prominently TRAK1 and BICDR1 – engage with Tyr^827^ through the originally proposed QxxH/Y motif (Fig. 7c; Fig. s7a-j)^3,6,7^. For most adaptors, the HBS1 tolerates markedly different sequence chemistries interacting with Tyr^827^ and the interaction is absent in two examples (HOOK3-A interaction with dynein-A2 and HOOK3-B interaction with dynein-A1). Conversely, almost all adaptors, except for NIN, displayed an acidic patch ∼15-20 residues after the Tyr^827^ interaction, which engages with Arg^759^. Previous studies have shown that charge reversal of the acidic patch is sufficient to inhibit dynein activation^5^, and HOOK3-A interacts with dynein-A2 through an acidic patch only (Fig. 6c). In all our adaptors, the HBS1 region is where the adaptor is best resolved, indicating that the interaction is a stable anchoring point for adaptor positioning in the dynein groove.

C-terminal to the HBS1 interaction, the trajectory of the adaptor coiled coil across the dynein groove is highly variable and significantly less stable. The sequences of the accessory HBS2 and HBS3 are not conserved, but both sites are consistently negatively charged, with limited exceptions (Fig. 7d). This is even true in TRAK1 where this region is flexibly bound. Conversely, the sites on dynein which recognise HBS2 and HBS3 are very well conserved (Fig. 7e-f) and positively charged (Fig. s7k). This organisation positions adaptors correctly for the contacts with dynactin’s pointed end, while accommodating a broad spectrum of sequences.

Earlier studies suggested that adaptors needed a continuous CC1 spanning from the HBS1 to the pointed end to activate dynein^3^. However, recent work has shown that adaptors with short coiled coils (JIP3) or broken ones (NUMA1) can also activate dynein motility efficiently^4,5^. Our results suggest that adaptors without breaks in this segment (e.g. BICD2 and BICDR1) are the exception rather than the rule. While the interactions of HBS1, HBS2, HBS3 and the binding to sites 2 and 3 of the pointed end complex involve coiled coils, continuity outside of these sites appears not to be necessary. NIN and RAB11FIP3 display short breaks after the HBS3 to allow the adaptor to correctly contact the dynactin pointed end. TRAK1 and NUMA1, instead, display breaks within the groove and a continuous coiled coil between HBS3 and the pointed end. Breaks also serve additional functions. In TRAK1, the break overlaps with the kinesin-1 binding region^19^. In Spindly, the coiled coil break is required for adaptor autoinhibition^7^. Overall, dynein-dynactin recognises a broad range of coiled coils through a set of conserved interfaces. Beyond these points, remarkable variation is possible and might allow for regulation of the interaction with dynein and, ultimately, motile output.

### The pointed end is a hub for adaptor interaction

Adaptor interactions at the dynactin pointed end also display considerable diversity. Four potential binding sites have been described previously^13^ (Fig. 7g). In our structures, interactions at sites 2, 3 and 4 are clearly visible (Fig. s8a). Our results allow us to better define site 2 interactions, and split the site in two, which we refer to as sites 2a and 2b (Fig. 7g; Fig. s8a-b). Site 2a is toward the top of the pointed end complex and binds HOOK3-B and NIN (Fig. 3e, 6g,i). Both adaptors interact with it via a pair of aspartates (Fig. 7h). Sequence alignments and AF2 predictions (Fig. s8c-e) suggest that other EF-hand adaptors NINL, RAB45 and RAB46 bind to the pointed end through a similar site 2a interaction. Other adaptors bind to site 2b, which is placed lower than 2a and involves a positively charged patch in the centre of p62 (Fig. 2e, 4e-f, 5i; Fig. s8b) and is formed by a flexible loop on p25^13^. FIP3, TRAK1 and BICD2 all bind both site 2b and site 3 (Fig. 2e, 4e-f, 5i). HOOK3-A binds only site 3, however (Fig. 6g). These interactions are less well resolved and more flexible than those with site 2a.

The existence of two productive binding modes at the pointed end raises intriguing functional questions. Site 2a interactions appear more stable than those at site 2b and 3, suggesting higher affinity. This agrees with kinetic data showing that NINL, which binds site 2a, assembles DDA complexes more rapidly than BICD2, a site 2b/3 adaptor^49^. Domain-swap experiments reinforce this idea: a BICD2–NINL chimera containing the BICD2 CC1 box and NINL coiled-coil, including the site 2-binding domain, formed complexes with higher affinity than wild-type BICD2, whereas the reciprocal chimera had reduced affinity compared with wild-type NINL^50^.

All adaptors additionally contact site 4 on the face of p25 via a Spindly motif or Spindly motif-like sequence. Based on this work, we can update the Spindly motif consensus sequence to JxxEΦ (with J = leucine or isoleucine, Φ = any small hydrophobic). The sole exception is NUMA1, where the central glutamate is substituted by a residue N-terminal of the Spindly motif^5^. Alignments identify potential JxxEΦ Spindly motifs in some adaptors for which structures are not yet known, such as HAP1, KASH5, and CDR2L adaptors (Fig. 7i). In contrast, the Spindly motif in RAB45 family adaptors was not obvious based on sequence alone. However AF2 did predict putative Spindly motif-like sequences in RAB45 and RAB46, albeit with lower confidence (Fig. 7i; Fig. s8f-g).

Surprisingly, an interaction with the pointed end is not absolutely required for adaptor function. In JIP3, the Spindly motif can be deleted with minimal effects on motility *in vitro*^4^. Furthermore, in the solved structure, the interaction with sites 2 and 3 is mediated by two coiled coil segments C-terminal of the Spindly motif and is also dispensable^20^. In FIP3, our classification shows the adaptor binds either the pointed end or via a different site to the actin subunit of the dynactin filament (Fig. 2e-f). FIP3 has been reported to contain binding sites for the cargo bound small G-proteins RAB11A and ARF5/6. Our structure suggests ARF5/6 binding could only occur with the adaptor in its actin subunit-bound position^32^. The ability of FIP3 and JIP3 to undock from the pointed end partially or fully thus provides a potential route for cargo binding to modulate the adaptor interaction with dynein-dynactin. Overall, our structures suggest that the dynactin pointed end is a central site of variability and that these sites may be involved in the regulation of the relative affinity of various adaptors for dynactin.

Taken together, our work defines a unifying framework for how activating adaptors engage and activate the dynein-dynactin machinery, while showing that this scaffold is implemented with far greater plasticity than previously appreciated. Rather than relying on a single conserved sequence, adaptors combine shared architectural anchors with variable coiled-coil organisation, alternative heavy-chain contacts and multiple pointed-end interactions, assembling motile complexes of different composition and likely different output. This flexibility provides a plausible molecular basis for how dynein is tuned for the diverse transport demands of human cells. More broadly, our findings establish a foundation for understanding how cargo binding, autoinhibition, and regulatory factors remodel adaptor behaviour, and offer a framework for interpreting disease-associated mutations that perturb transport. These principles should guide future efforts to link adaptor identity to transport function in cells.

## Acknowledgements

We thank A. Rico Montaña for help with DDT data collection. We thank G. Manigrasso for protein used in the DDB structure. We thank the MRC Laboratory of Molecular Biology Electron Microscopy Facility for access and support of electron microscopy sample preparation and data collection; and J. Grimmett, T. Darling, and I. Clayson for providing scientific computing resources. We acknowledge Diamond Light Source for access and support of the cryo-EM facilities at the UK national electron Bio-Imaging Centre (eBIC), proposal BI31336. This work was supported by UK Research and Innovation Horizon Europe Funding Guarantee MSCA Postdoctoral Fellowship (101105484) to E.A.d’A., an EMBO Postdoctoral Fellowship (ALTF 334-2020) to S.C., a Sir Henry Wellcome Postdoctoral Fellowship to F.A.A. (218653/Z/19/Z), Boehringer Ingelheim Fonds PhD fellowship to L.M., and a Wellcome Trust Investigator award (227434/Z/23/Z) and MRC funding grant (MC_UP_A025_1011) to A.P.C. For the purpose of open access, the MRC Laboratory of Molecular Biology has applied a CC BY public copyright licence to any Author Accepted Manuscript version arising.

## Author Contributions

E.A.d’A. and A.P.C. conceived the study and designed the experiments. S.C. developed the SUBFLOW software. E.A.d’A. solved the structures of dynein-dynactin in complex with FIP3, NIN, BICD2 and HOOK3. E.A.d’A. and S.C. solved the structure of dynein-dynactin in complex with TRAK1. E.A.d’A. and F.A.A. performed data acquisition and cryo-EM grid preparation for the HOOK3 structure. E.A.d’A. and L.M. performed data acquisition and cryo-EM grid preparation for the NIN structure. E.A.d’A. built and refined all models. E.A.d’A. and A.P.C. wrote the manuscript, and all authors read and revised the manuscript.

## Methods

### Constructs

FIP3 FL, FIP3 (1-719) and Rabin8 were generated by inserting the two constructs into a pAceBac1 plasmid for insect cell expression, with an N-terminal 2xStrep-PSc tag. Rab11 Q70L mutant, which is GTP-locked was cloned into a pET28a plasmid with an N-terminal 2xStrep-SNAP tag.

NIN (1-693) was generated by inserting the construct into a pAceBac1 plasmid for insect cell expression, with an N-terminal 2xStrep-EGFP-TEV tag.

TRAK1 (1-421) was cloned into a pAceBac1 plasmid for insect cell expression with an N-terminal SNAP and a C-terminal PSc-Strep tag. Two constructs of *M. musculus* BICD2 were used: MBP-TEV-BICD2 (1-658)-His_6_ and 2xStrep-PSc-BICD2 (1-400), both inserted into a pAceBac1 plasmid for insect cell expression.

Two constructs of HOOK3 were used: HOOK3 (1-522) was generated by inserting the construct into a pAceBac1 plasmid with a C-terminal SNAP-PSc-Strep-His_6_ tag, and the FHF construct used was previously published^47^.

A previously published dynein *phi* mutant construct (DYNC1H1 R1567E and K1610E) was used to recombinantly express dynein in Sf9 cells^9^. Previously published ZZ-LIS1 and ZZ-LIS1-SNAP in pFastBac vector were used for expression of LIS1 in insect cells^4^.

### Protein expression

All insect cell expressions were carried out in Sf9 cells. Sf9 cultures were infected with 1% (v/v) P_2_ virus and grown for 72 h at 27 °C before harvesting by centrifugation and snap-freezing. Expression of Rab11 Q70L mutant was performed in SoluBL21 cells, induced at an OD_600_ of 0.6 with 1 mM IPTG and cultured overnight at 18 °C with shaking. Cells were harvested by centrifugation and snap-frozen.

### Protein purification

All adaptors except TRAK1 and the FHF complex were purified in GF150 buffer (25 mM HEPES pH 7.2, 150 mM KCl, 1 mM MgCl_2_, 1 mM DTT). Pellets from 1 l Sf9 or bacterial culture were resuspended in 30 ml GF150 buffer with 1 mM PMSF and cOmplete protease inhibitor cocktail (Roche). Lysis was performed by sonication, and the lysate clarified by high-speed centrifugation (100,000 *g*) for 50 min at 2 °C. The supernatant was collected and filtered through a 0.8 µm filter. We used two-step purification with a first affinity step, followed by gel filtration. For Strep-tagged protein, we performed the affinity step loading the filtered lysate on a 1 mL StrepTrap HP column (Cytiva), followed by a wash and elution with GF150 with 5 mM desthiobiotin. For His-tagged protein, we performed the affinity step loading the filtered lysate on a 5 mL HisTrap HP column (Cytiva), followed by a wash and elution with GF150 with 300 mM imidazole.

In both cases, the fractions of the peak were determined by SDS-PAGE, pooled, concentrated, and loaded on a Sepharose S200 Increase column (Cytiva) for gel filtration. The fractions containing the protein of interest were determined by SDS-PAGE, and the quality of the protein was confirmed by measuring protein mass using mass photometry (Refeyn MP).

TRAK1 was purified from 1 l Sf9 culture in a two-step procedure consisting of Strep affinity purification followed by protease cleavage and size-exclusion chromatography. Cell pellets were resuspended in 40 ml lysis buffer (50 mM HEPES pH 7.4, 150 mM KAc, 2 mM MgSO4, 1 mM EGTA, 10% (v/v) glycerol, 1 mM DTT, 0.5% Triton X-100, 2 mM PMSF, and EDTA-free protease inhibitor cocktail), incubated for 15 min at 4 °C, and clarified by ultracentrifugation at 70,000 rpm for 45 min in a Type 70 Ti rotor. The supernatant was incubated with 2 ml Strep-Tactin resin (IBA) for 2 h at 4 °C, after which the resin was washed extensively with lysis buffer and then with PreScission cleavage buffer (30 mM HEPES pH 7.4, 150 mM NaCl, 1 mM EDTA, 1 mM DTT).

Bound TRAK1 was released by overnight cleavage with PreScission protease at 4 °C. The eluate was concentrated and loaded onto a Superose 6 Increase 10/300 column equilibrated in GF150 buffer. Peak fractions were pooled, concentrated, glycerol was added to a final concentration of 10%, and the protein was aliquoted, snap frozen in liquid nitrogen, and stored at -80 °C.

Purification of the FHF complex was performed as previously described^47^. Briefly, pellets from 2 l culture were thawed and resuspended in 45 ml lysis buffer (50 mM HEPES pH 7.2, 150 mM NaCl, 10% glycerol, 1 mM DTT) supplemented with 2 mM PMSF and EDTA-free protease inhibitors. Cells were lysed by douncing (20–25 strokes) on ice and clarified by ultracentrifugation at 504,000g for 45 min at 4 °C. The lysate was applied to 4 ml pre-equilibrated Strep-Tactin Sepharose beads, with the flow-through reapplied to maximise binding. Beads were washed with ∼450 ml lysis buffer followed by 200 ml PreScission (Psc) buffer. Beads were resuspended in Psc buffer, split into two tubes, and incubated with PreScission protease overnight at 4 °C with rotation. Protein was subjected to size exclusion chromatography on a Superose 6 10/300 column, and peak fractions were pooled and concentrated. Glycerol was added to 10%, and the protein was aliquoted and stored.

Purification of dynein *phi* mutant was performed as previously described^4,8^. Briefly, pellet from 1 l Sf9 culture was resuspended in 30 ml lysis buffer (50 mM HEPES pH 7.2, 100 mM KCl, 10% (v/v) glycerol, 1 mM DTT, 0.1 mM ATP), supplemented with one tablet cOmplete protease inhibitor cocktail. Lysis was performed by douncing with 10 cycles. Lysate was clarified at 60,000 rpm in a Type 70 Ti rotor (Beckman Coulter) for 45 mins at 2 °C. The clarified lysate was decanted and incubated for at least 4 h with 3 ml pre-equilibrated Immunoglobulin G (IgG) Sepharose 6 Fast Flow beads (Cytiva). The beads were transferred to a gravity column and washed with at least 30 CV of lysis and TEV buffer (50 mM TRIS-HCl pH 7.4, 150 mM KAc, 2 mM MgAc_2_, 1 mM EGTA, 10% (v/v) glycerol, 1 mM DTT, 0.1 mM ATP). Beads were transferred to two 2 ml Eppendorf tubes filled completely with TEV buffer. 200 µg TEV protease was added to each tube, and the sample was incubated overnight at 4 °C on a roller. The supernatant, containing the cleaved dynein, was eluted with a gravity column, and the beads were washed to a final eluate volume of approximately 7 ml. The protein was collected and concentrated no higher than 2 mg/ml, then loaded on a TSKgel G4000_SWXL_ column (TOSOH Bioscience) pre-equilibrated with GF150 buffer with 5 mM DTT and 0.1 mM ATP. Peak fractions were pooled and concentrated to a final concentration of around mg/ml. Glycerol was added to a concentration in the final freezing buffer of 10%. The purified protein was aliquoted and stored in 3 µl aliquots at -80 °C.

Purification of LIS1 was performed essentially as previously reported^4^. A pellet from 1 l Sf9 culture was resuspended in 30 ml lysis buffer B (50 mM TRIS-HCl pH 8, 250 mM KAc, 2 mM MgAc_2_, 1 mM EGTA, 10% (v/v) glycerol, 0.1 mM ATP, 1 mM DTT) supplemented with cOmplete protease inhibitor cocktail. Cells were lysed by douncing with 10 cycles. Lysate was clarified at 60,000 rpm in a Type 70 Ti rotor for 45 min at 4 °C. The clarified lysate was incubated with 3 ml pre-equilibrated IgG Sepharose Fast Flow beads for 4 h at 4 °C on a roller. The beads were then applied to a gravity column and washed with at least 30 CV lysis buffer B, after which they were transferred to two 2 ml Eppendorf centrifuge tubes. 200 µg TEV protease was added to each tube and incubated overnight at 4 °C for tag cleavage. The supernatant, containing the cleaved protein, was eluted with a gravity column, and the beads were washed to a final eluate volume of approximately 7 ml. The protein was concentrated to 3-5 mg/ml and loaded onto a Superdex S200 Increase column (Cytiva), pre-equilibrated in GF150 buffer. Peak fractions were pooled and concentrated to around 3-4 mg/ml. Aliquots were flash frozen and stored at -80 °C until use.

Purification of native dynactin from pig brain was performed as previously published^4^.

### Cryo-EM sample preparation

Cryo-EM samples were prepared essentially as described previously^5^. Briefly, microtubules were prepared using GMPCPP (most datasets) or paclitaxel (FIP3 datasets 1-4, HOOK3 dataset 1) as a stabiliser, and mixed with a pre-formed complex of dynein-dynactin-adaptor-LIS1. We did not see differences between complexes in presence of paclitaxel or GMPCPP, but we do note that GMPCPP-stabilised microtubules are more regular and therefore tend to yield better subtraction.

For GMPCPP-stabilised microtubules, the microtubule mixture was prepared incubating porcine tubulin at a concentration of 6 µM in BRB80 buffer (80 mM PIPES pH 6.8, 1 mM MgCl_2_, 1 mM EGTA) with 1 mM GMPCPP for 1h at 37 °C. Microtubules were pelleted by centrifugation at 20,000 *g* for 8 minutes, resuspended in BRB80, and depolymerised for 30 minutes on ice. A second polymerisation round was started by addition of 1 mM GMPCPP and incubation at 37 °C for 1h, followed by another round of centrifugation and resuspension in BRB80 to a final concentration of 0.3 mg/ml, as measured by Bradford assay (BioRad).

For paclitaxel-stabilised microtubules, 6 µM tubulin was polymerised for 1.5 h at 37 °C in BRB80 (80 mM PIPES pH 6.8, 1 mM MgCl_2_, 1 mM EGTA, 1 mM DTT) with 10% DMSO and 1 mM Mg.GTP. The sample was diluted 1:5 with BRB80-T buffer (BRB80 supplemented with 20 µM paclitaxel), and centrifuged for 8 minutes at 21,000g at room temperature. The resulting pellet was resuspended in 80 µl BRB80-T buffer and pelleted again for 8 minutes at 21,000g at room temperature. Microtubules were resuspended in BRB80-T to a final concentration of 0.6 mg/ml, as measured by Bradford assay.

Dynein-dynactin-LIS1-adaptor complexes were formed by mixing the purified proteins at a 1:2:50:50 ratio in GF150 buffer (25 mM HEPES pH 7.2, 150 mM KCl, 1 mM MgCl_2_, 1 mM DTT), with dynein at a concentration of 0.07 µM, to a final volume of 10 µl, with or without LIS1, and incubating 30 minutes on ice.

The cryo-EM sample was prepared by mixing 9 µl of complex mixture, 9 µl of microtubule mixture, and 5 µl of buffer S1 (25 mM HEPES pH 7.2, 5 mM MgSO_4_, 1 mM EGTA, 2 mM DTT, 25 mM AMPPNP) and incubating 15 minutes at room temperature. This allowed dynein-dynactin-adaptor-LIS1 complexes to bind tightly to microtubules. The microtubules were pelleted by centrifugation at 20,000 *g* for 8 minutes, after which the pellet was dried and resuspended in S2 buffer (25 mM HEPES pH 7.2, 35 mM KCl, 5 mM MgSO_4_, 1 mM EGTA, 1 mM DTT, 5.3 mM AMPPNP, 0.01% IGEPAL (Millipore-Sigma)). For paclitaxel-stabilised microtubules, S1 and S2 buffers also included 52 and 20 µM paclitaxel respectively.

The final resuspension volume varied between samples and always ranged between 70-100 µl. 4 µl of sample was applied to a glow-discharged Quantifoil R2/2 mesh 300 gold grid (Quantifoil) and blotted at 22 °C and 100% humidity in a Vitrobot IV (ThermoFisher), with 45 s wait time, blot force -15 N, and blotting times ranging between 1.5 and 2.5 s.

### Cryo-EM data acquisition

Grids were screened using a Glacios or Titan Krios microscope (Thermo Fisher Scientific) at the LMB EM facility. Data acquisition was performed on multiple microscopes: at the LMB EM facility on a Titan Krios equipped with a K3 detector with an energy filter (20 eV slit, Gatan), at the LMB EM facility on a Titan Krios equipped with a Falcon 4 detector without energy filter, at the Diamond eBIC facility on an FEI Titan Krios equipped with a K3 detector with an energy filter (20 eV slit, Gatan), and at the Diamond eBIC facility on an FEI Titan Krios equipped with a Falcon 4 and Selectris X energy filter. Detailed information on all data collections is available in supplementary Table 1.

### Cryo-EM image processing

Image processing was carried out in Subflow (https://github.com/sami-chaaban/subflow), an internally developed graphical user interface that enables on-the-fly monitoring of incoming files and orchestrates the microtubule-subtraction workflow together with downstream RELION and cryoSPARC processing. Incoming movies were linked into the working directory and, where applicable, EER movies were converted to TIFF before preprocessing. Global motion correction, dose-weighting, coordinate import, particle extraction, 3D refinement and polishing were performed in RELION^51,52^, and CTF parameters were estimated with CTFFIND4^53^. Microtubules were first detected with crYOLO in standard particle mode, rather than filament mode^54^. These coordinates were then processed with the multi-curve-fitting (MCF) procedure, which assigns helical-tube IDs and resamples positions along each microtubule^24^. The resampled coordinates were split into shorter fragments, typically 10 points per segment, before lattice subtraction to reduce artefacts from microtubule bending. Subtraction was performed with the tubulin-lattice-subtraction workflow^24^ wrapped by Subflow, using an automatically scaled mask. Where beneficial, a second round of microtubule detection, MCF, splitting and subtraction was performed to improve subtraction quality and increase particle density. Candidate dynein-dynactin-adaptor particles were then picked from the subtracted micrographs with crYOLO in standard particle mode.

Particles picked from the subtracted micrographs were imported into RELION for extraction. We aimed to use an extraction box corresponding to ∼880 Å. For the first extraction, particles were binned to ∼2.5 Å per pixel for the BICD2, HOOK3, FIP3 and NIN datasets 2 and 3, ∼3 Å per pixel for NIN dataset 1, and ∼4.4 Å per pixel for TRAK1. Extracted particles were imported into cryoSPARC for heterogeneous refinement^55^ using eight reference volumes: six representing common contaminants, one a dynein-dynactin-BICD2 complex with two dynein dimers, and one the corresponding one-dynein-dimer complex. In all datasets except BICD2, the one-dynein-dimer class degraded during heterogeneous refinement and was not pursued further, although the presence of a very small number of such particles in the other datasets cannot be excluded. To exploit the stochasticity of cryoSPARC heterogeneous refinement for particle recovery, in select cases the refinement was repeated three or five times on the same particle stack, and the resulting particle lists were combined with duplicates removed. Selected particles then underwent non-uniform refinement in cryoSPARC^55^ and were converted back to RELION.star format using pyem^56^. In RELION, binned particles underwent a single round of 3D refinement, generally with BLUSH regularization enabled^51^, which facilitated generation of tighter focused masks. Refined particles were then polished and re-binned to a standard pixel size of approximately 1.3 Å per pixel for BICD2, NIN, FIP3 and HOOK3, and approximately 1.7 Å per pixel for TRAK1. At this stage, multiple datasets were combined and subjected to a first round of joint refinement.

The overall resolutions reached at this step were under 4 Å, but the estimation was greatly inflated by the dynactin filament which is extremely well-resolved. We leveraged this for CTF parameter estimation, which we performed on signal-subtracted particles where only the density for the highest-resolution parts of the structure was kept, namely the dynactin barbed end and the first few subunits of ARP1 filament. Particles were reverted to the full particle, previous alignments restored, and 3D-refinement was performed again, yielding substantially improved maps. After this step, particle subtraction was performed on the regions of interest. In all cases, broadly similar masks were used to refine the HBS1, dynein-A tail, and pointed end regions. Local classifications were performed in RELION with BLUSH off. A sample pre-processing pipeline to the point of local refinement can be seen in Fig. s1. Data collection and image processing statistics can be found in supplementary tables 1-2.

Composite maps were generated by placing locally refined maps into the consensus map for each complex using UCSF ChimeraX. The maps were then aligned with respect to the consensus using the “fit in map” function. The maps were merged using the volume max function and then filtered to 10 Å. In the case of the DDB one-dynein complex, the consensus map was filtered to 10 Å and used for further model building.

### Model building and refinement

Model building was performed in ISOLDE^57^ and Coot^58^. Refinement was performed using PHENIX^59^. Starting models were docked into density using UCSF ChimeraX^60^. We used either Protein Data Bank (PDB) models as starting models or AF2-Multimer^61^ predictions. AF2 predictions were generated using a local installation of Colabfold 5^62^ using MMSeq^63^ for homology searches.

To build full complex models, we first used AF2 predictions of the HBS1-dynein heavy chain interface to establish the coiled-coil register within the dynein–dynactin groove. Additional structural landmarks, including pointed-end contacts, pseudo-EF-hand positions in NIN, coiled-coil breaks, and focused high-resolution maps of the HBS1 interaction, were used to validate this register assignment. Linearised AF2 models of each adaptor coiled coil were then docked into the cryo-EM density and flexibly refined in ISOLDE under restraining potentials. AF2 predictions of adaptor-pointed end contacts guided placement of the Spindly motif. In the case of HOOK3, as the density at the HOOK domains was poor and did not allow us to confidently resolve the domain, AF predictions for the domain were docked into the density based on the position of the CC1a which immediately followed them, and refined in ISOLDE under high restraints.

Dynactin and the dynein tails were modelled using PDB 8PTK as a starting point. Whereas dynactin remained rigid across reconstructions, the dynein tails, and particularly dynein-A1, showed substantial variability. The 8PTK model was therefore docked jointly with the pre-fitted adaptor, followed by global flexible fitting in ISOLDE using strong restraints. Regions not supported by density were removed in the final model. The models were then automatically refined in PHENIX.

The one-dynein model of the DDB complex was built based on the two-dynein model, where dynein-B was deleted and the adaptor and dynein-A was flexibly fit into the density in ISOLDE with strong restraints. The model was then manually refined in Coot and automatically refined in PHENIX.

To build models of the HBS1 interaction with dynein-A2 and B1, and HBS2/3 interactions with dynein-A1 and A2, we docked the full-complex model into the focused density, and deleted residues outside the density. The resulting models were manually refined in Coot and automatically refined in PHENIX. The densities were sufficient to assign registry accurately, but in all cases, except HOOK3, we were aided by high-confidence AF2 predictions of the HBS1-dynein interaction. In the case of TRAK1, density was too poor to position the acidic patch confidently and we were helped by distance constraints from the Tyr^827^ interaction, as well as the relevant AF2 prediction. For HOOK3 interaction with dynein-A1, we could use AF2 as a model. For the interactions with dynein-A2 and B1, we used the density to place the coiled coil and then refined in coot with restraints.

To model adaptor–pointed-end interactions, we docked AF2 predictions generated using the four pointed-end subunits (p25, p27, p62 and ARP11, *Sus scrofa* homologues) together with two copies of a short adaptor segment selected based on our full-complex models and sequence homology around the Spindly motif. For HOOK3, AF2 was run with multiple fragment combinations, and the prediction that best satisfied the density and geometric constraints was chosen as the starting model. This model was then docked into the map, and segments unsupported by density were removed.

Models of the site 2 + site 4 interactions (NIN, HOOK3) showed excellent agreement with the cryo-EM density from the outset and required only manual refinement and trimming in Coot. By contrast, models of the site 3 + site 4 interactions were substantially more flexible, with the region of adaptor interacting with site 3 displaying mobility. The adaptor was in this case flexibly fit into the experimental density with restraints. The models were then automatically refined using PHENIX, with restraints.

## Data availability

Cryo-EM structures have been deposited to PDB and EMDB and will be available upon publication. The constructs used in this study are available from the corresponding author upon request. Raw EM data will be made available by the corresponding author upon request.

## Code availability

The SUBFLOW code used for micrograph preprocessing is available in GitHub (https://github.com/sami-chaaban/subflow).

