## Supplementary figures for "Structural basis of dynein interaction with diverse activating adaptors"

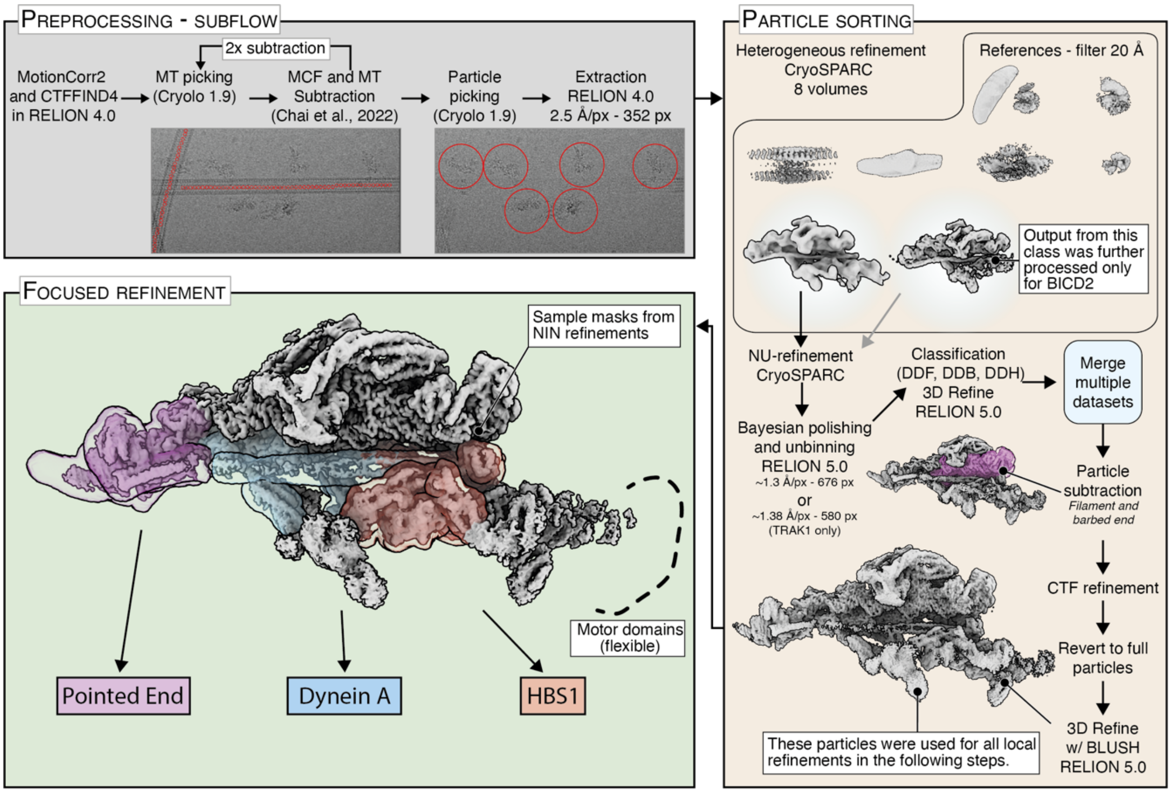


**Supplementary Figure 1: Processing pipeline for SPA of DDA complexes on microtubules.** A graphical user interface (Subflow) was developed to handle preprocessing steps (see Methods). For TRAK1, box and pixel size for pre-processing were 200 px and 4.4 Å respectively.

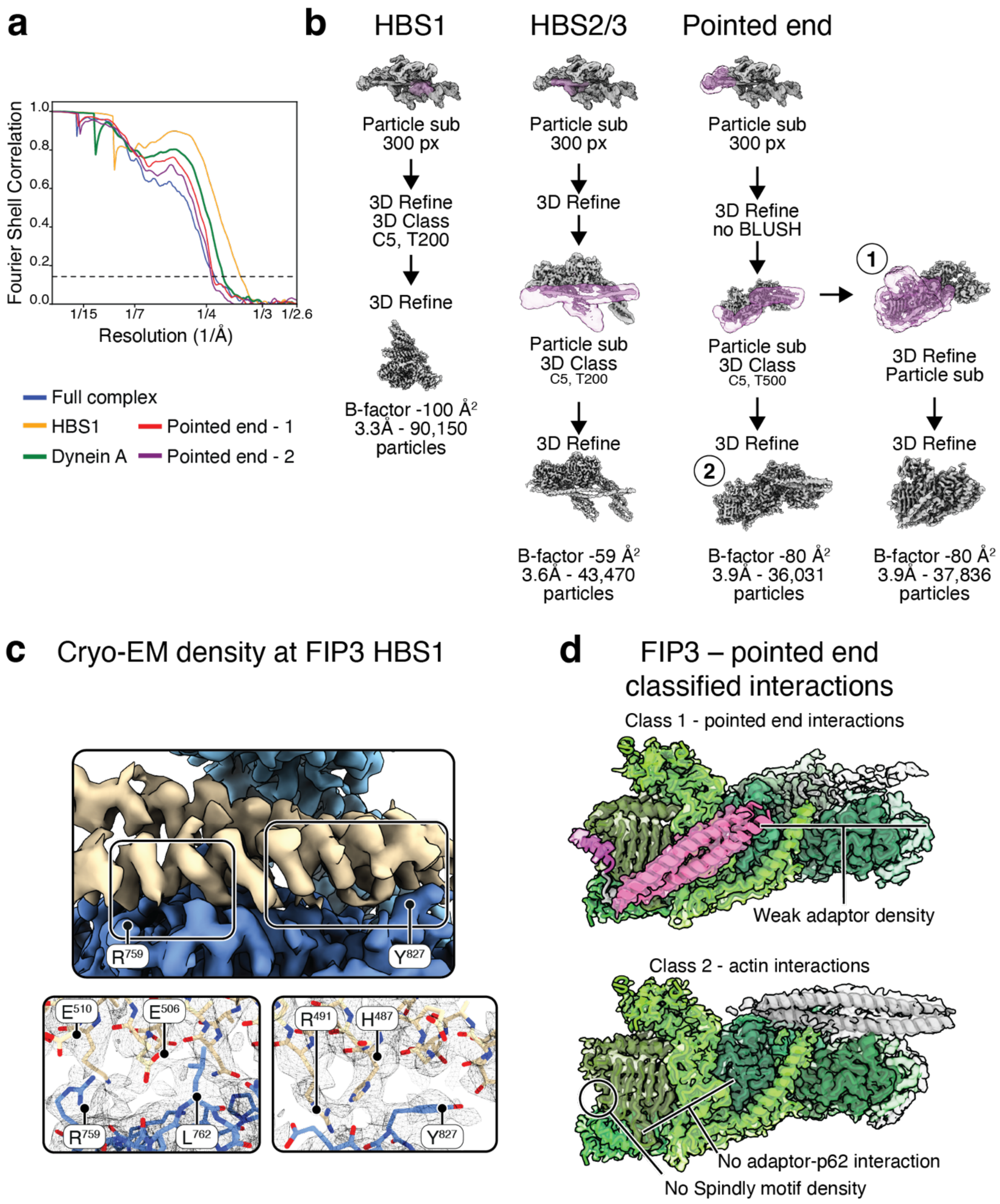


**Supplementary Figure 2: Supplementary analysis of the dynein-dynactin-RAB11FIP3 complex.** **a**, FSC plot showing gold-standard Fourier shell correlation. The dotted line shows the 0.143 cut-off. **b**, Processing pipelines for DDF subcomplexes with refinement information. **c**, density map at the HBS1 of RAB11FIP3, showing the position of the Arg$^{759}$ and Tyr$^{827}$-mediated interactions and the model fit at these two sites. **d**, Density map of the pointed end of RAB11FIP3 in classes 1 and 2, showing the interaction between adaptor and pointed end in class 1, and the interaction between adaptor and actin in class 2.

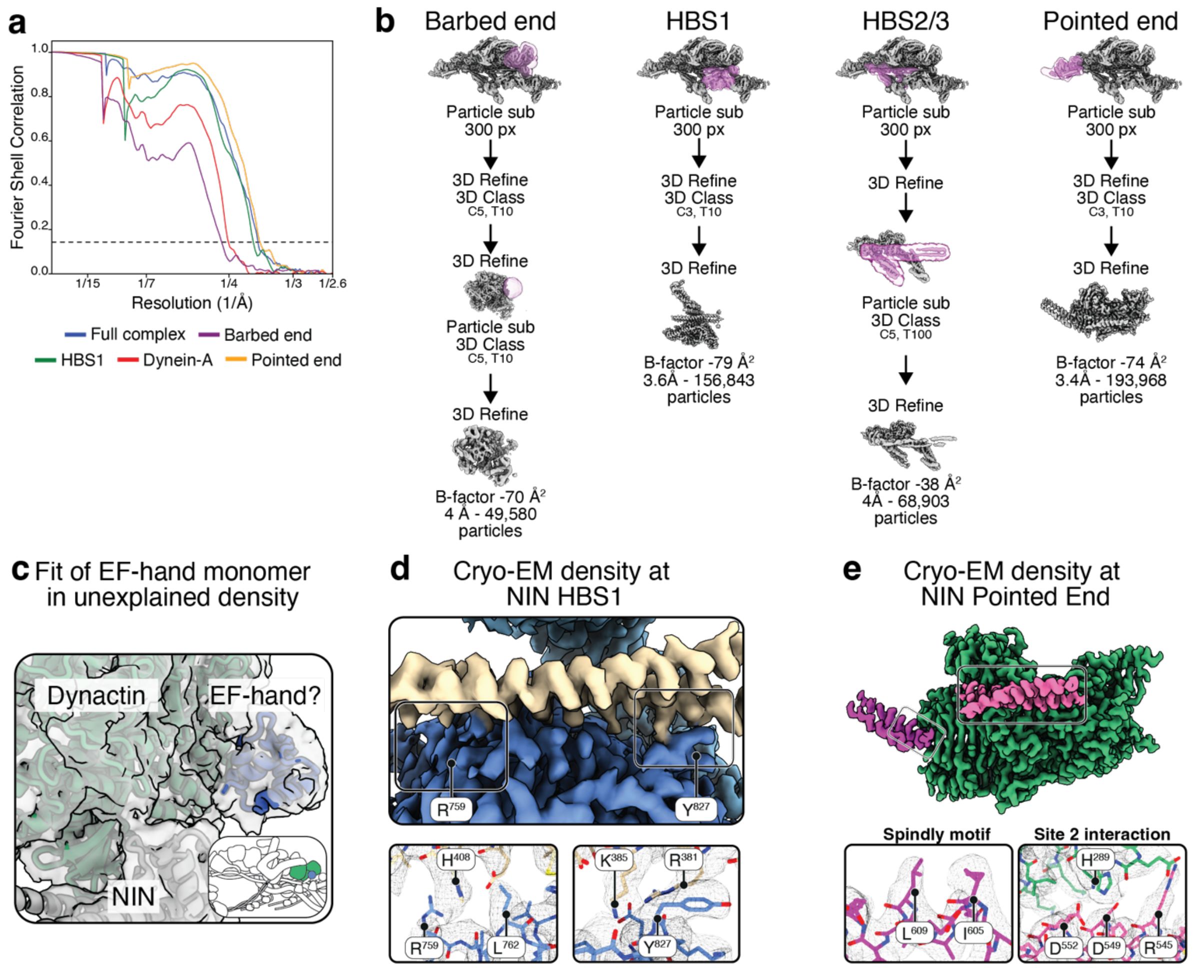


**Supplementary Figure 3: Supplementary analysis of the dynein-dynactin-NIN complex. a**, FSC plot showing gold-standard Fourier shell correlation. The dotted line shows the 0.143 cut-off. **b**, Processing pipeline for DDN subcomplexes with refinement information. **c**, Low-resolution fit depicting EF-hand pair 2 in NIN in the unexplained density off the dynactin pointed end. EF-hand pair 1 could also fit (not shown). **d**, Density map at the HBS1 of NIN, showing the position of the $Arg^{759}$ and $Tyr^{827}$-mediated interactions and the residue fit at these two sites. **e**, Density map at the pointed end, showing the position of the Spindly motif and site 2a interactions and the residue fit at these two sites.

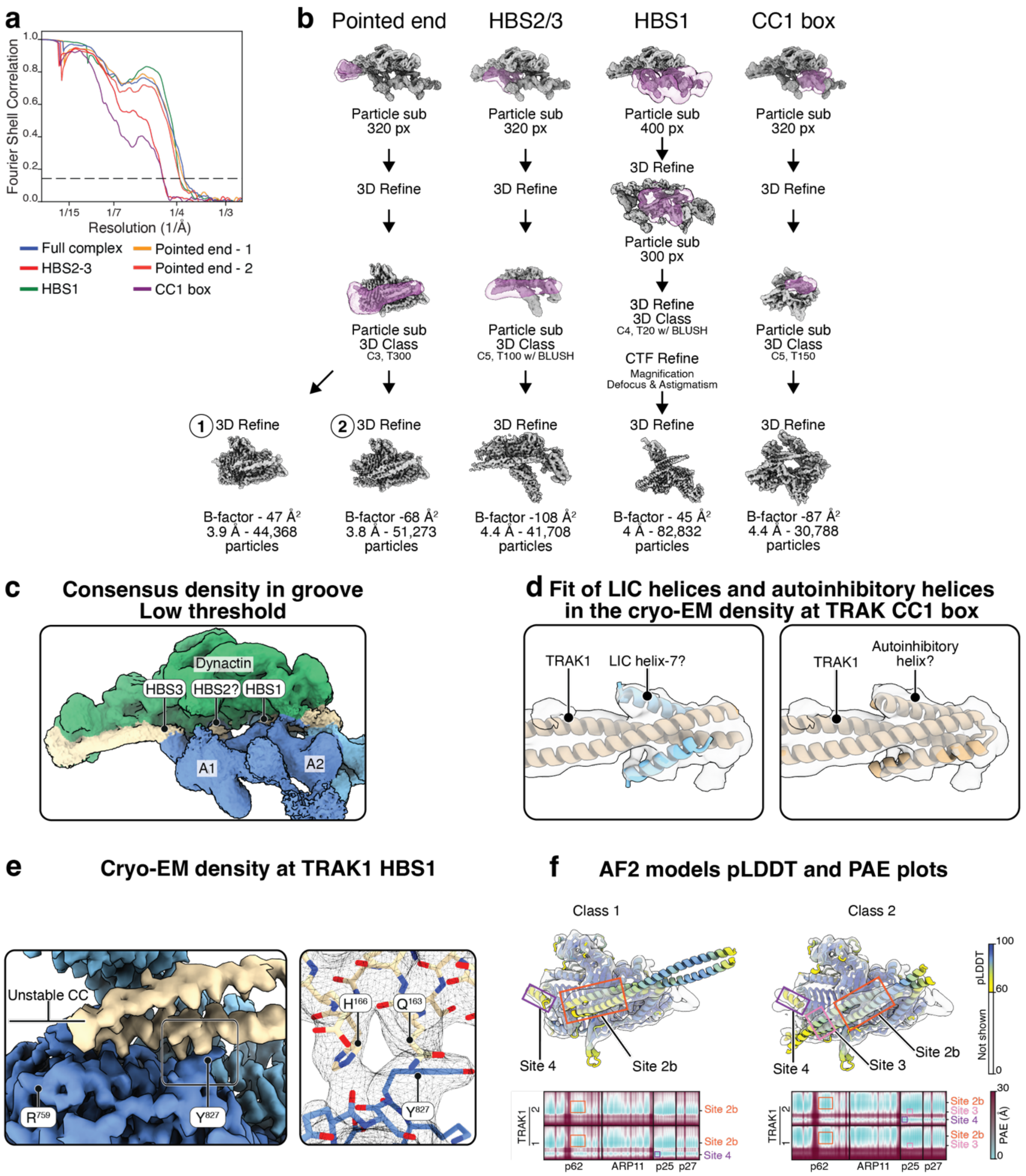


**Supplementary Figure 4: Supplementary analysis of the dynein-dynactin-TRAK1 complex**. **a**, FSC plot showing gold-standard Fourier shell correlation. The dotted line shows the 0.143 cut-off. **b**, Processing pipelines for DDT subcomplexes with refinement information. **c**, Filtered consensus map for DDT complex at the HBS2-3 sites, with low threshold, shows unexplained density at the HBS2. **d**, Fit of an AF2 prediction of the LIC-CC1 box interaction in TRAK1, with autoinhibitory helices unengaged, and of an AF2 prediction of the TRAK1 N-terminal alone, with autoinhibitory helices engaged in our CC1 density. **e**, Density map at the HBS1 of TRAK1, showing the position of the Tyr$^{827}$-mediated interactions and the residue fit at the site. **f**, Top: fit of AF2 models in filtered densities for the interaction between pointed end and TRAK1 (position 1, left; position 2, right). Coloured by pLDDT. Flexible linkers with pLDDT < 60 were hidden to assist visualisation. Bottom: relevant PAE plot for the two chains of TRAK1 against the subunits of the pointed end. Coloured squares correlate the position of each interaction site in the model with the PAE plot and show low PAE values at interaction sites.

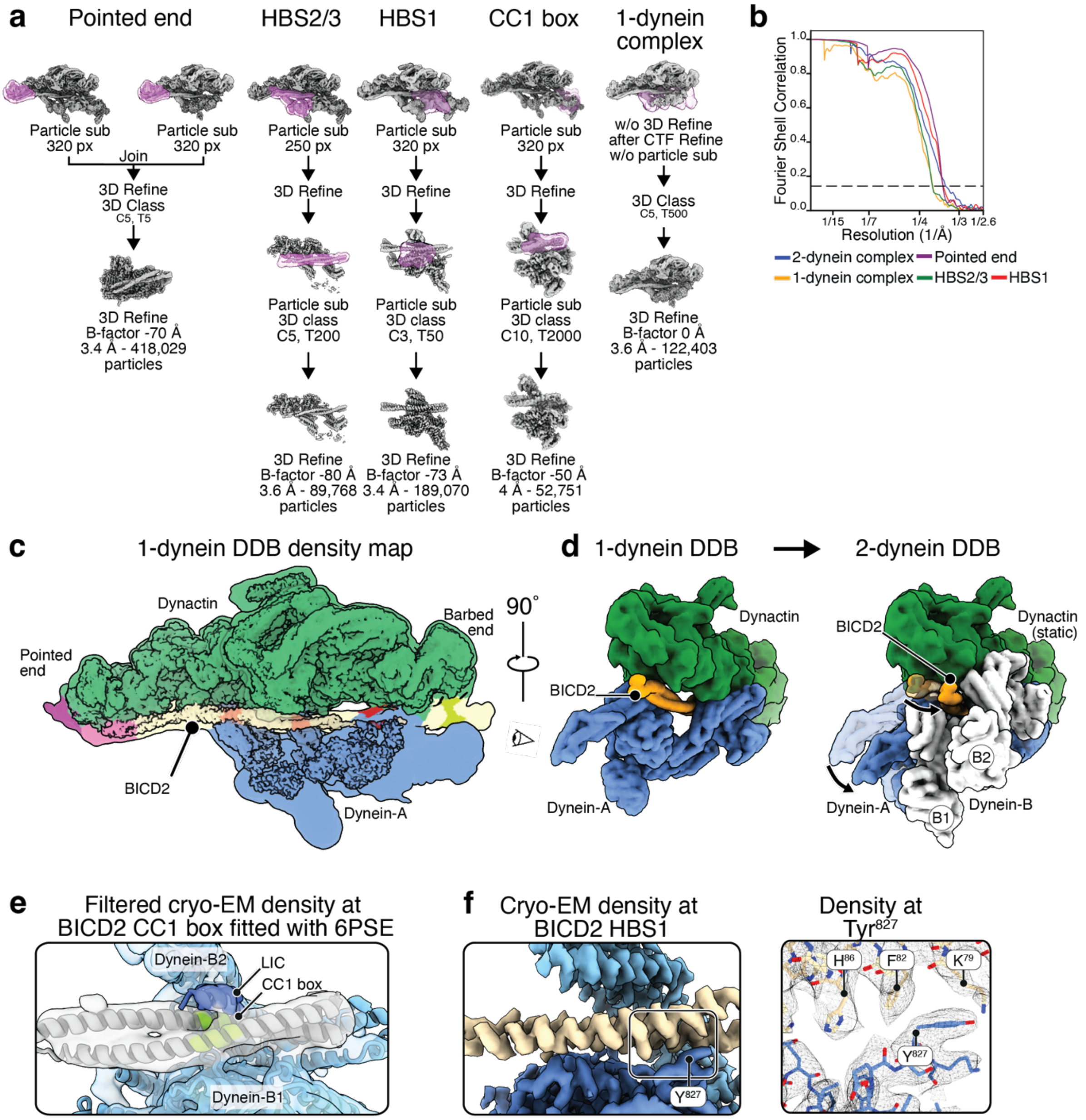


**Supplementary Figure 5: Supplementary analysis of the dynein-dynactin-BICD2 complex**. **a**, Processing pipelines for DDB subcomplexes with refinement information. **b**, FSC plot showing gold-standard Fourier shell correlation. The dotted line shows the 0.143 cut-off. **c**, Consensus density map for the dynein-dynactin-BICD2 complex when only a single dynein dimer is present. The dynein motors are flexible relative to the adaptor-interaction module and are therefore not resolved here. **d**, Model depicting the movement of the adaptor and dynein-A relative to dynactin when switching from the one-dynein to two-dynein complex. Adaptor in orange, dynein-A in blue, dynein-B in white. Arrows indicate the direction of the swing between the 1-dynein and the 2-dynein conformations. **e**, Rigid-body fit of the crystal structure of LIC-bound BICD2 (PDB: 6PSE) in the density at the CC1 box shows lack of a second bound LIC. Dynein-B position is based on the composite structure. **f**, Density map at the HBS1 of BICD2, showing the position of the $Tyr^{827}$-mediated interaction and the residue fit at this site.

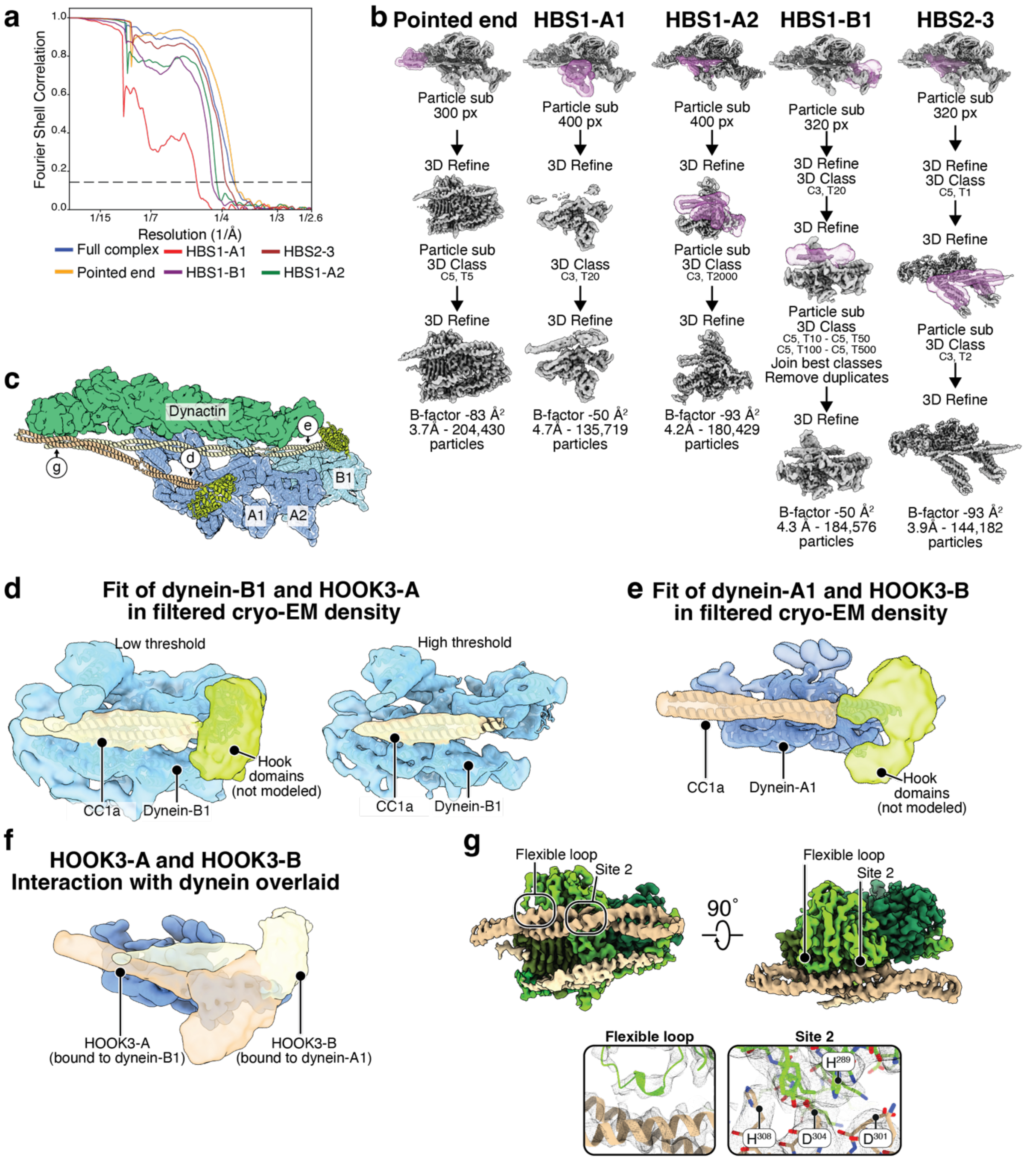


**Supplementary Figure 6: Supplementary analysis of the dynein-dynactin-HOOK3 complex.** **a**, FSC plot showing gold-standard Fourier shell correlation. The dotted line shows the 0.143 cut-off. **b**, Processing pipelines for DDH subcomplexes with refinement information. **c**, Space-filling representation of the position of main and accessory HOOK3 adaptors in the DDH complex. Subunits of the dynactin shoulder have been hidden for visibility reasons. **d**, Fit of the HOOK3-A and dynein-B1 model in the cryo-EM density, at two different thresholds showing density for the HOOK domain and fit for the CC1a segment. **e**, Fit of the HOOK3-B and dynein-A1 model in the cryo-EM density, showing the unmodeled density for the HOOK domain. **f**, Filtered densities for HOOK3-A and HOOK3-B with dynein-B1 and dynein-A2 overlaid show that the coiled coil is differently oriented. **g**, EM density at the pointed end interaction with HOOK3-A and B and fit of our model in the density. Insets show density and model fit at specific sites.

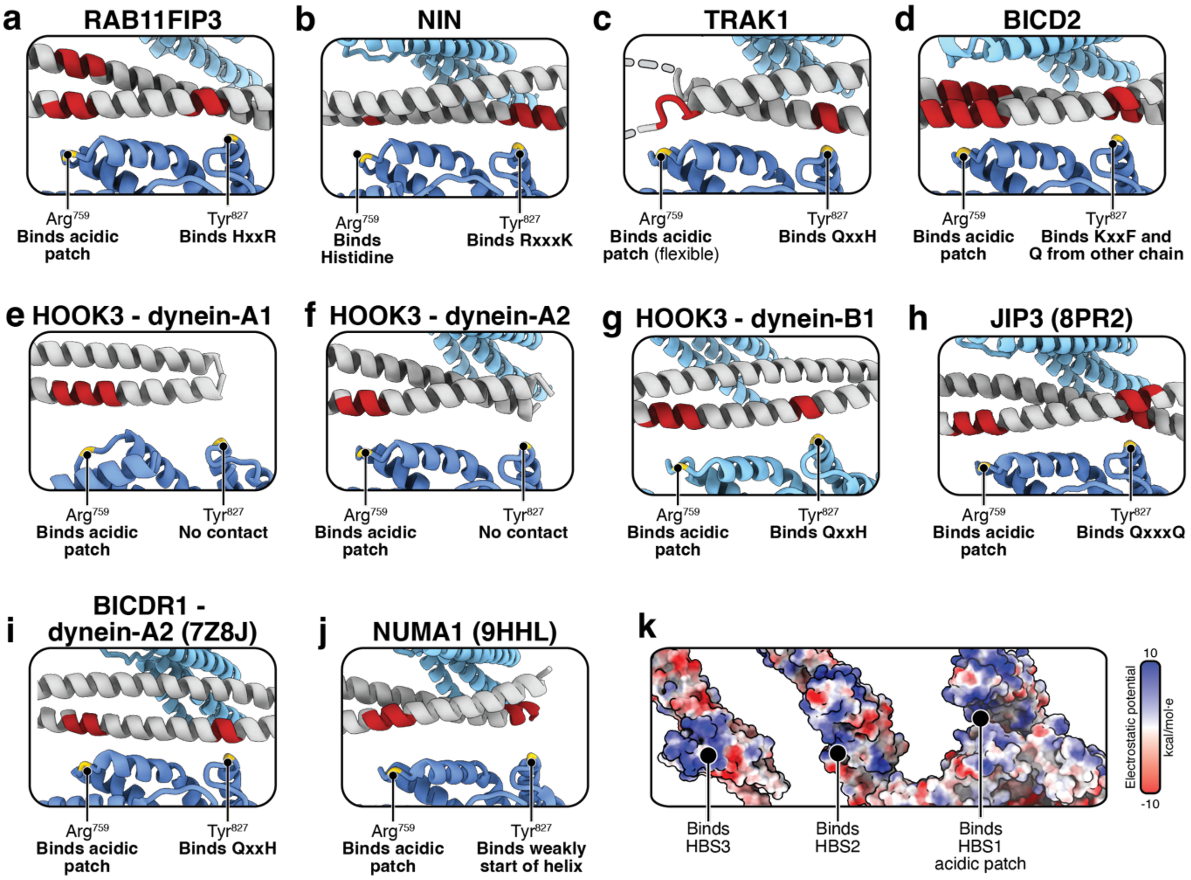


**Supplementary Figure 7: Interactions at the HBS1-3. a-j**, HBS1-like interactions between dynein heavy chain and adaptors RAB11FIP3 (**a**), NIN (**b**), TRAK1 (**c**), BICD2 (**d**), HOOK3-B interaction with dynein-A1 (**e**), HOOK3-A interaction with dynein-A2 (**f**), HOOK3-A interaction with dynein-B1 (**g**), JIP3 (**h**), BICDR1-A interaction with dynein-A2 (**i**), NUMA1 (**j**). **k**, Dynein surface representation in the groove, coloured by coulombic potential. The model is a superimposition of the dynein models for the HBS2-3 and HBS1 regions in BICD2.

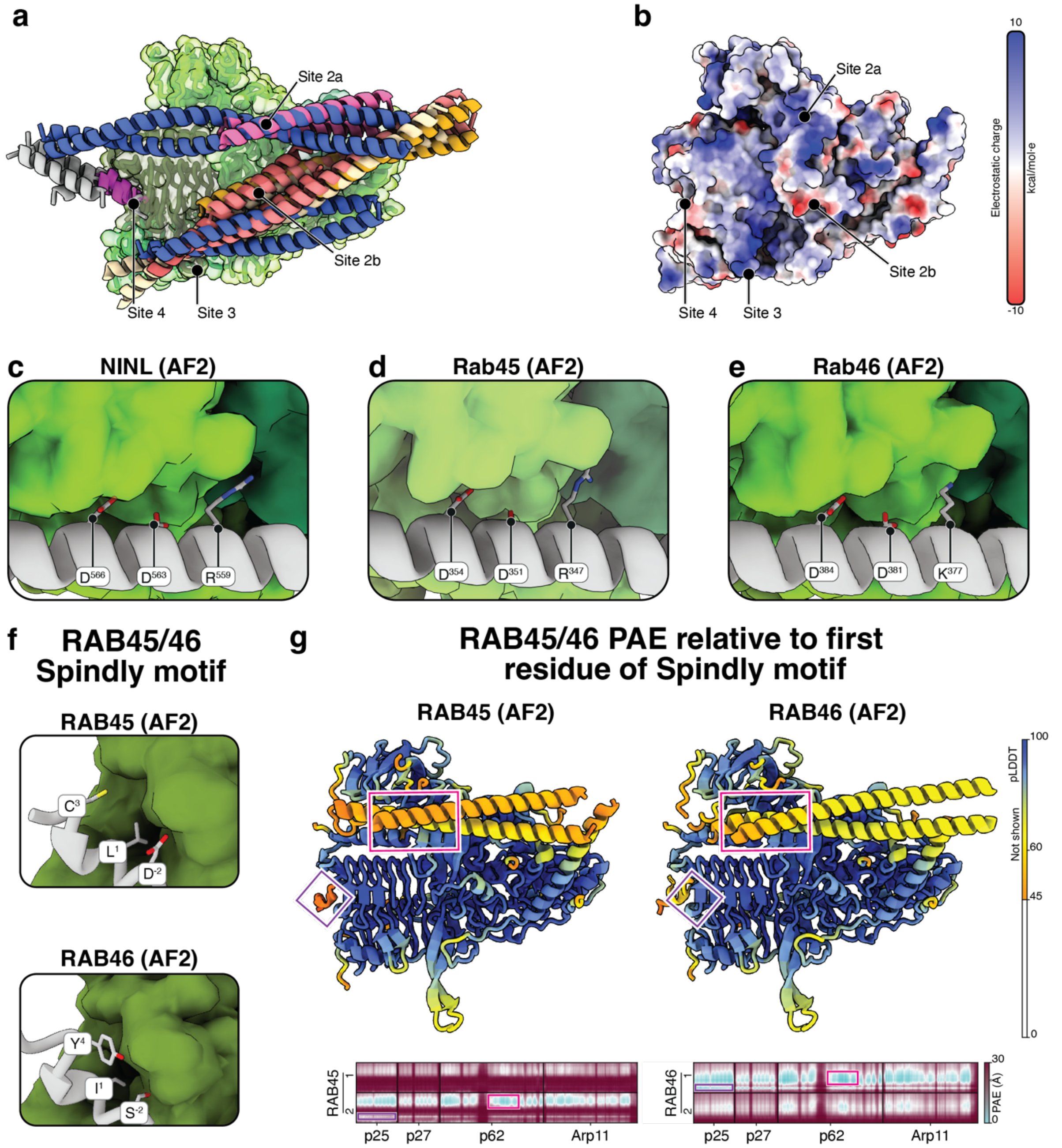


**Supplementary Figure 8: Pointed End - adaptor interactions. a**, All adaptors in this study overlaid onto the pointed end for the NIN model, showing variety of interactions with the pointed end. All Spindly motifs and CC2 coloured in magenta and silver. Blue, HOOK3. Orange, TRAK1 class 1. Beige, TRAK1 class 2. Pink, NIN. Coral, BICD2. Red, RAB11FIP3. **b**, Structure of the dynactin pointed end coloured by coulombic potential. **c-e**, AF2 predictions of site 2 binding of adaptors RAB45 (**c**), NINL (**d**), and RAB46 (**e**). **f**, AF2 predictions of unusual Spindly motifs in RAB45 and RAB46. **g**, AF2 model of RAB45 and RAB46 bound to the pointed end, coloured by PAE relative to the first residue of the Spindly motif showing the accuracy of the prediction. Relevant PAE plots below, with the Spindly motif (magenta) and site 2a (pink) interactions within boxes.
