## Supplementary table 1 for "Structural basis of dynein interaction with diverse activating adaptors"

**Supplementary table 1**: Cryo-EM refinement statistics

| *FIP3 complexes* | *HBS1* | *HBS2-3* | *Pointed end (1)* | *Pointed end (1) – large map* | *Pointed end (2)* | *Consensus* | *Composite* |
|---:|---|---|---|---|---|---|---|
| ***EMDB ID*** | 57637 | 57648 | 57639 | 57634 | 57641 | 57636 | 57643 |
| ***PDB ID*** | 30DK | 30DM | 30DN | - | 30DP | - | 30DN |
| *Particle images* | 90,150 | 43,470 | 37,836 | 37,836 | 36,031 | 159,084 | - |
| *Symmetry* | C1 | C1 | C1 | C1 | C1 | C1 | C1 |
| *Map Resolution* | 3.3 Å | 3.6 Å | 3.9 Å | 3.9 Å | 3.9 Å | 3.8 Å | Filtered to 10 Å |
| *FSC threshold* | 0.143 | 0.143 | 0.143 | 0.143 | 0.143 | 0.143 | - |
| *Sharpening B-factor (Å$^2$)* | -100 | -59 | -80 | -80 | -80 | -71 | - |
| ***Model Refinement*** | | | | | | | |
| *Model Resolution (Å)* | 3.4 | 3.8 | 3.9 | - | 3.9 | - | - |
| *FSC threshold* | 0.5 | 0.5 | 0.5 | - | 0.5 | - | - |
| ***Model composition*** | | | | | | | |
| *Non-hydrogen atoms* | 8,013 | 7,927 | 8,552 | - | 13,755 | - | 69,030 |
| *Protein residues* | 1,011 | 1,589 | 1,189 | - | 1,954 | - | 13,948 |
| *Ligands* | - | ADP:2, Mg:2 | ATP:1, Zn:3 | - | ATP:1, Zn: 3, ADP:2, Mg: 2 | - | - |
| ***Mean B-factors (Å$^2$)*** | | | | | | | |
| *Protein* | 55.93 | 63.64 | 79.32 | - | 30.08 | - | 639.04 |
| *Ligand* | - | 13.67 | 68.87 | - | 14.72 | - | - |
| ***RMSD deviations*** | | | | | | | |
| *Bond lengths (Å)* | 0.006 | 0.005 | 0.003 | - | 0.004 | - | 0.002 |
| *Bond angles (°)* | 0.845 | 1.090 | 0.659 | - | 0.809 | - | 0.435 |
| ***Validation*** | | | | | | | |
| *Clashscore* | 6.51 | 0.35 | 6.51 | - | 5.75 | - | 0.61 |
| *MolProbity score* | 1.43 | 0.88 | 1.37 | - | 1.66 | - | 0.83 |
| *Rotamers outliers (%)* | 0.83 | - | 0.12 | - | 0.59 | | - |
| ***Ramachandran plot*** | | | | | | | |
| *Favored (%)* | 97.67 | 96.25 | 97.94 | - | 95.07 | | 97.32 |
| *Allowed (%)* | 2.33 | 3.75 | 2.06 | - | 4.93 | | 2.67 |
| *Outliers (%)* | - | - | - | - | - | | 0.01 |

| ***Ninein complexes*** | ***HBS1*** | ***HBS2-3*** | ***Pointed end*** | ***Barbed end*** | ***Consensus*** | ***Composite*** |
|---|---|---|---|---|---|---|
| ***EMDB ID*** | 57647 | 57646 | 57648 | 57645 | 57644 | 57649 |
| ***PDB ID*** | 30DT | 30DS | 30DU | - | - | 30DV |
| *Particle images* | 193,968 | 68,903 | 193,968 | 49,580 | 308,720 | - |
| *Symmetry* | C1 | C1 | C1 | C1 | C1 | C1 |
| *Map Resolution* | 3.6 Å | 4 Å | 3.4 Å | 4 Å | 3.5 Å | Filtered to 10 Å |
| *FSC threshold* | 0.143 | 0.143 | 0.143 | 0.143 | 0.143 | - |
| *Sharpening B-factor (Å$^2$)* | -79 | -38 | -74 | -40 | -50 | - |
| ***Model Refinement*** | | | | | | |
| *Model Resolution (Å)* | 3.6 | 4.1 | 3.5 | - | - | - |
| *FSC threshold* | 0.5 | 0.5 | 0.5 | - | - | - |
| ***Model composition*** | | | | | | |
| *Non-hydrogen atoms* | 10,411 | 7,867 | 9,261 | - | - | 65,164 |
| *Protein residues* | 1,420 | 1,577 | 1,220 | - | - | 13,167 |
| *Ligands* | - | ADP:2, Mg:2 | ATP:1, Zn:3 | - | - | - |
| ***Mean B-factors (Å$^2$)*** | | | | | | |
| *Protein* | 34.47 | 106.42 | 55.59 | - | - | 601.68 |
| *Ligand* | - | 54.17 | 42.86 | - | - | - |
| ***RMSD deviations*** | | | | | | |
| *Bond lengths (Å)* | 0.004 | 0.003 | 0.005 | - | - | 0.004 |
| *Bond angles (°)* | 0.679 | 0.552 | 0.820 | - | - | 1.009 |
| ***Validation*** | | | | | | |
| *Clashscore* | 3.83 | 0.53 | 2.63 | - | - | 0.63 |
| *MolProbity score* | 1.33 | 1.15 | 1.39 | - | - | 0.81 |
| *Rotamers outliers (%)* | - | - | - | - | - | - |
| ***Ramachandran plot*** | | | | | | |
| *Favored (%)* | 97.08 | 92.58 | 95.14 | - | - | 97.52 |
| *Allowed (%)* | 2.92 | 7.42 | 4.86 | - | - | 2.47 |
| *Outliers (%)* | - | - | - | - | - | 0.02 |

| *TRAK1 complexes* | *HBS1* | *HBS2-3* | *Pointed end (1)* | *Pointed end (2)* | *CC1 box* | *Consensus* | *Composite* |
|---|---|---|---|---|---|---|---|
| ***EMDB ID*** | 57661 | 57658 | 57659 | 57660 | 57650 | 57651 | 57662 |
| ***PDB ID*** | 30EE | 30ED | - | - | - | - | 30EF |
| *Particle images* | 82,832 | 41,708 | 44,368 | 51,273 | 30,788 | 129,749 | - |
| *Symmetry* | C1 | C1 | C1 | C1 | C1 | C1 | C1 |
| *Map Resolution* | 4 Å | 4.4 Å | 3.9 Å | 3.8 Å | 4.4 Å | 3.8 Å | Filtered to 10 Å |
| *FSC threshold* | 0.143 | 0.143 | 0.143 | 0.143 | 0.143 | 0.143 | - |
| *Sharpening B-factor (Å$^2$)* | -45 | -38 | -47 | -68 | -87 | -30 | - |
| ***Model Refinement*** | | | | | | | |
| *Model Resolution (Å)* | 4 | 4.4 | - | - | - | - | - |
| *FSC threshold* | 0.5 | 0.5 | - | - | - | - | - |
| ***Model composition*** | | | | | | | |
| *Non-hydrogen atoms* | 9,737 | 6,537 | - | - | - | - | 64,652 |
| *Protein residues* | 1,295 | 1,320 | - | - | - | - | 13,064 |
| *Ligands* | - | - | - | - | - | - | - |
| ***Mean B-factors (Å$^2$)*** | | | | | | | |
| *Protein* | 61.12 | 67.24 | - | - | - | - | 652.27 |
| *Ligand* | - | - | - | - | - | - | - |
| ***RMSD deviations*** | | | | | | | |
| *Bond lengths (Å)* | 0.005 | 0.002 | - | - | - | - | 0.003 |
| *Bond angles (˚)* | 0.784 | 0.444 | - | - | - | - | 0.595 |
| ***Validation*** | | | | | | | |
| *Clashscore* | 5.49 | 0.11 | - | - | - | - | 0.35 |
| *MolProbity score* | 1.46 | 0.54 | - | - | - | - | 0.83 |
| *Rotamers outliers (%)* | 0.86 | - | - | - | - | - | - |
| ***Ramachandran plot*** | | | | | | | |
| *Favored (%)* | 97.11 | 98.14 | - | - | - | - | 96.78 |
| *Allowed (%)* | 2.89 | 1.86 | - | - | - | - | 3.19 |
| *Outliers (%)* | - | - | - | - | - | - | 0.02 |

| ***BICD2 complexes*** | ***HBS1*** | ***HBS2-3*** | ***Pointed end*** | ***CC1 box*** | ***Consensus*** | ***One-dynein consensus*** | ***Composite*** |
|---|---|---|---|---|---|---|---|
| ***EMDB ID*** | 57632 | 57633 | 57689 | 57631 | 57688 | 57716 | 57697 |
| ***PDB ID*** | 30DE | 30DH | 30FD | - | - | 30FU | 30FL |
| *Particle images* | 189,070 | 89,768 | 418,029 | 52,751 | 433,095 | 122,403 | - |
| *Symmetry* | C1 | C1 | C1 | C1 | C1 | C1 | C1 |
| *Map Resolution* | 3.4 Å | 3.6 Å | 3.4 Å | 4.0 Å | 3.3 Å | Filtered to 5 Å | Filtered to 10 Å |
| *FSC threshold* | 0.143 | 0.143 | 0.143 | 0.143 | 0.143 | - | - |
| *Sharpening B-factor (Å$^2$)* | -73 | -80 | -47 | 0 | -69 | - | - |
| ***Model Refinement*** | | | | | | | |
| *Model Resolution (Å)* | 3.4 | 3.7 | 3.4 | - | - | - | - |
| *FSC threshold* | 0.5 | 0.5 | 0.5 | - | - | - | - |
| ***Model composition*** | | | | | | | |
| *Non-hydrogen atoms* | 9,300 | 10,847 | 8,993 | - | - | 49,871 | 67,878 |
| *Protein residues* | 1,181 | 1,396 | 1,246 | - | - | 10,082 | 13,715 |
| *Ligands* | - | ADP:2, Mg:2 | ATP:1, Zn:3 | - | - | - | - |
| ***Mean B-factors (Å$^2$)*** | | | | | | | |
| *Protein* | 58.05 | 53.09 | 54.28 | - | - | 123.28 | 642.11 |
| *Ligand* | - | 54.17 | 23.79 | - | - | - | - |
| ***RMSD deviations*** | | | | | | | |
| *Bond lengths (Å)* | 0.005 | 0.009 | 0.008 | - | - | 0.003 | 0.003 |
| *Bond angles (°)* | 0.756 | 1.170 | 0.973 | - | - | 0.572 | 0.423 |
| ***Validation*** | | | | | | | |
| *Clashscore* | 5.73 | 7.48 | 3.62 | - | - | 0.94 | 1.25 |
| *MolProbity score* | 1.31 | 1.70 | 1.46 | - | - | 1.02 | 0.97 |
| *Rotamers outliers (%)* | 0.62 | 0.28 | 0.47 | - | - | - | - |
| ***Ramachandran plot*** | | | | | | | |
| *Favored (%)* | 98.19 | 95.78 | 95.58 | - | - | 96.47 | 97.36 |
| *Allowed (%)* | 1.81 | 4.22 | 4.42 | - | - | 3.53 | 2.62 |
| *Outliers (%)* | - | - | - | - | - | 0.01 | 0.01 |

| HOOK3 complexes | HBS1-A1 | HBS1-A2 | HBS1-B1 | HBS2-3 | Pointed end | Consensus | Composite |
|---|---|---|---|---|---|---|---|
| **EMDB ID** | 57671 | 57690 | 57722 | 47720 | 57664 | 57703 | 57724 |
| **PDB ID** | 30EM | 30FE | 30FX | 30FW | 30EI | - | 30FY |
| *Particle images* | 135,719 | 180,429 | 418,029 | 122,403 | 52,751 | 426,971 | - |
| *Symmetry* | C1 | C1 | C1 | C1 | C1 | C1 | C1 |
| *Map Resolution* | 4.7 Å | 4.2 Å | 4.3 Å | 3.9 Å | 3.7 Å | 3.8 Å | Filtered to 10 Å |
| *FSC threshold* | 0.143 | 0.143 | 0.143 | 0.143 | 0.143 | 0.143 | - |
| *Sharpening B-factor (Å$^2$)* | -50 | -93 | -50 | -93 | -83 | -80 | - |
| **Model Refinement** | | | | | | | |
| *Model Resolution (Å)* | 4.8 | 4.2 | 4.3 | 4 | 3.8 | - | - |
| *FSC threshold* | 0.5 | 0.5 | 0.5 | 0.5 | 0.5 | - | - |
| **Model composition** | | | | | | | |
| *Non-hydrogen atoms* | 5,230 | 8,536 | 7,702 | 8,681 | 10,377 | - | 74,438 |
| *Protein residues* | 1,054 | 1,099 | 1,554 | 1,753 | 1,397 | - | 15,037 |
| *Ligands* | - | - | - | - | Zn:3, ATP:1 | - | - |
| **Mean B-factors (Å$^2$)** | | | | | | | |
| *Protein* | 171.85 | 58.92 | 91.54 | 63.78 | 70.60 | - | 609.57 |
| *Ligand* | - | - | - | - | 55.28 | - | - |
| **RMSD deviations** | | | | | | | |
| *Bond lengths (Å)* | 0.004 | 0.008 | 0.003 | 0.002 | 0.006 | - | 0.002 |
| *Bond angles (°)* | 0.501 | 0.987 | 0.568 | 0.588 | 0.759 | - | 0.393 |
| **Validation** | | | | | | | |
| *Clashscore* | 1.99 | 2.40 | 0 | 0.08 | 3.79 | - | 0.92 |
| *MolProbity score* | 1.3 | 1.36 | 0.6 | 0.85 | 1.33 | - | 0.96 |
| *Rotamers outliers (%)* | - | 0.23 | - | - | 0.76 | - | 0 |
| **Ramachandran plot** | | | | | | | |
| *Favored (%)* | 95.16 | 95.09 | 97.52 | 95.45 | 97.08 | - | 96.97 |
| *Allowed (%)* | 4.84 | 4.91 | 2.48 | 4.55 | 2.92 | - | 3.03 |
| *Outliers (%)* | - | - | - | - | - | - | 0.01 |
