## Supplementary table 2 for "Structural basis of dynein interaction with diverse activating adaptors"

**Supplementary table 2:** Cryo-EM data collection statistics

| *FIP3 Dataset* | *1* | *2* | *3* | *4* | *5* | *6* | *7* |
|---|---|---|---|---|---|---|---|
| *Sample* | DDL + FIP3 (1-719) | DDL + FIP3 | DDL + FIP3 | DDL + FIP3 | DDL + FIP3 + RAB11 + RABIN8 | DDL + FIP3 + RAB11 + RABIN8 | DDL + FIP3 + RAB11 + RABIN8 |
| *Facility* | LMB | LMB | LMB | LMB | LMB | LMB | eBIC |
| *Voltage (kV)* | 300 | 300 | 300 | 300 | 300 | 300 | 300 |
| *Camera* | K3 + GIF | K3 + GIF | Falcon 4 | K3 + GIF | K3 + GIF | Falcon 4 | Falcon 4 + EF |
| *Pixel size (Å)* | 0.93 | 0.93 | 1.08 | 0.93 | 1.059 | 1.1 | 1.185 |
| *Electron exposure* | 50 $e^-/Å^2$ | 50 $e^-/Å^2$ | 50 $e^-/Å^2$ | 50 $e^-/Å^2$ | 50 $e^-/Å^2$ | 50 $e^-/Å^2$ | 50 $e^-/Å^2$ |
| *Defocus range* | -1.2 to -2.2 | -1.2 to -2.2 | -1.2 to -2.2 | -1.2 to -2.2 | -1.2 to -2.2 | -1.2 to -2.2 | -1.2 to -2.2 |
| *Total micrographs* | 11,242 | 7,266 | 13,084 | 5,328 | 12,307 | 8,810 | 21,221 |
| *Initial particles* | 415,101 | 252,182 | 147,235 | 158,888 | 390,944 | 194,522 | 740,785 |
| *Cleaned particles* | 25,702 | 14,942 | 19,534 | 20,014 | 17,734 | 18,180 | 76,512 |

| *NIN Dataset* | *1* | *2* | *3* |
|---|---|---|---|
| *Sample* | DDL + NIN (1-693) | DDL + NIN (1-693) | DDL + NIN (1-693) |
| *Facility* | LMB | LMB | LMB |
| *Voltage (kV)* | 300 | 300 | 300 |
| *Camera* | Falcon 4 | Falcon 4 | Falcon 4 |
| *Pixel size (Å)* | 1.1 | 1.1 | 1.1 |
| *Electron exposure* | 40 $e^-/Å^2$ | 40 $e^-/Å^2$ | 40 $e^-/Å^2$ |
| *Defocus range* | -1.2 to -2.2 | -1.2 to -2.2 | -1.2 to -2.2 |
| *Total micrographs* | 28,671 | 47,202 | 52,404 |
| *Initial particles* | 249,806 | 1,197,768 | 1,246,484 |
| *Cleaned particles* | 37,800 | 131,919 | 139,001 |

| *TRAK1 Dataset* | *1* | *2* | *3* | *4* | *5* | *6* | *7* | *8* |
|---|---|---|---|---|---|---|---|---|
| *Sample* | DDL + TRAK (1-421) | DDL + TRAK (1-421) | DDL + TRAK (1-421) + KIF5B | DDL + TRAK (1-421) | DDL + TRAK (1-421) | DDL + TRAK (1-421) + KIF5B + MAP7 | DDL + TRAK (1-421) + KIF5B + MAP7 | DDL + TRAK (1-421) + KIF5B + MAP7 |
| *Facility* | LMB | LMB | LMB | LMB | eBIC | LMB | LMB | LMB |
| *Voltage (kV)* | 300 | 300 | 300 | 300 | 300 | 300 | 300 | 300 |
| *Camera* | Falcon 4 | Falcon 4 + EF | K3 + GIF | Falcon 4 | K3 + GIF | Falcon 4 | Falcon 4 | Falcon 4 |
| *Pixel size (Å)* | 1.1 | 1.222 | 1.11 | 1.1 | 1.058 | 1.1 | 1.1 | 1.1 |

| | | | | | | | | |
|---|---|---|---|---|---|---|---|---|
| *Electron exposure* | 50 e⁻/Å² | 50 e⁻/Å² | 50 e⁻/Å² | 50 e⁻/Å² | 50 e⁻/Å² | 50 e⁻/Å² | 50 e⁻/Å² | 50 e⁻/Å² |
| *Defocus range* | -1.2 to -2.8 | -1.0 to -2.4 | -1.4 to -2.6 | -1.2 to -2.4 | -1.2 to -2.2 | -1.2 to -2.2 | -1.2 to -2.2 | -1.2 to -2.2 |
| *Total micrographs* | 16,786 | 5,312 | 8,719 | 31,128 | 19,129 | 6,477 | 7,763 | 9,060 |
| *Initial particles* | 162,514 | 95,727 | 326,335 | 414,512 | 521,880 | 94,586 | 96,144 | 123,232 |
| *Cleaned particles* | 7,808 | 8,032 | 20,673 | 37,102 | 41,489 | 5,012 | 8,143 | 6,334 |

| *BICD2 Dataset* | *1* | *2* | *3* | *4* | *5* |
|---|---|---|---|---|---|
| *Sample* | DDL + BICD2 1-658 | DDL + BICD2 1-400 | DD + BICD2 1-400 | DD + BICD2 1-400 | DDL + BICD2 1-400 |
| *Facility* | LMB | LMB | LMB | eBIC | eBIC |
| *Voltage (kV)* | 300 | 300 | 300 | 300 | 300 |
| *Camera* | Falcon 4 | K3 + EF | Falcon 4 | K3 + EF | K3 + EF |
| *Pixel size (Å)* | 1.1 | 1.164 | 1.1 | 1.072 | 1.072 |
| *Electron exposure* | 50 e⁻/Å² | 50 e⁻/Å² | 50 e⁻/Å² | 50 e⁻/Å² | 50 e⁻/Å² |
| *Defocus range* | -1.2 to -2.2 | -1.2 to -2.2 | -1.2 to -2.2 | -1.2 to -2.2 | -1.2 to -2.2 |
| *Total micrographs* | 27,428 | 26,244 | 30,862 | 38,054 | 48,509 |
| *Initial particles* | 670,156 | 840,551 | 631,206 | 987,482 | 1,607,993 |
| *Cleaned particles (one-dynein class)* | 23,973 | 28,055 | 49,316 | 71,132 | 97,494 |
| *Cleaned particles (two-dynein class)* | 41,802 | 42,164 | 106,064 | 106,736 | 154,487 |

| *HOOK3 Dataset* | *1* | *2* | *3* | *4* | *5* | *6* |
|---|---|---|---|---|---|---|
| *Sample* | DDL + FHF + Kif1C | DDL + HOOK3 (1-522) | DDL + HOOK3 (1-522) | DDL + HOOK3 (1-522) | DDL + HOOK3 (1-522) | DDL + HOOK3 (1-522) |
| *Facility* | LMB | LMB | LMB | LMB | LMB | LMB |
| *Voltage (kV)* | 300 | 300 | 300 | 300 | 300 | 300 |
| *Camera* | K3 + GIF | Falcon 4 | Falcon 4 | Falcon 4 | K3 + GIF | Falcon 4 |
| *Pixel size (Å)* | 1.11 | 1.1 | 1.1 | 1.1 | 1.12 | 1.1 |
| *Electron exposure* | 50 e⁻/Å² | 40 e⁻/Å² | 40 e⁻/Å² | 40 e⁻/Å² | 40 e⁻/Å² | 40 e⁻/Å² |
| *Defocus range* | -1.2 to -2.2 | -1.2 to -2.2 | -1.2 to -2.2 | -1.2 to -2.2 | -1.2 to -2.2 | -1.2 to -2.2 |
| *Total micrographs* | 16,318 | 32,637 | 10,424 | 9,782 | 43,299 | 10,866 |
| *Initial particles* | 910,390 | 1,029,620 | 332,437 | 260,068 | 1,427,236 | 331,072 |
| *Cleaned particles* | 132,890 | 147,693 | 55,531 | 26,494 | 245,718 | 34,993 |
